# Prion-like characteristics of Hepatitis E virus ORF3 protein are associated with virus release and pathogenesis

**DOI:** 10.1101/2023.01.13.523929

**Authors:** Yajing Wang, Houlu Tian, Na Shi, Chunyan Wu, Yaowei Huang, Yongling Yang, Qiang Ding, Xu Zheng, Qin Zhao, Mick F. Tuite, Hongying Chen, Yuchen Nan

## Abstract

Hepatitis E virus (HEV) is a common causative agent of acute hepatitis. Due to the shortage of efficient in vitro model, the viral assembly and release processes are still poorly understood. In this study, we found that ORF3, an HEV structural protein showed prion-like properties. The prion domain (PrD) of yeast prion Sup35 could be functionally replaced by HEV-ORF3 and its candidate PrD (cPrD). A single amino acid substitution in the cPrD reduced the aggregation propensity of ORF3 and blocked the function of ORF3 in enhancing the stability of microtubules in HEV-infected cells, thus blocking virion release from the infected cells and resulting in reduced HEV pathogenicity in Mongolian gerbils. These data suggest that HEV-ORF3 is a novel functional prion-like protein which assembles into a filamentous structure for virion release, supporting the hypothesis that the self-propagating properties of prion proteins are widely exploited as epigenetic information carriers in nature.

## 1. Introduction

Hepatitis E virus (HEV) is a quasi-enveloped, single-stranded positive-sense RNA virus. It is categorized within an expanding family of *Hepeviridae*^1,2^, which contains several zoonotic, anthropotropic and animal-restricted HEV species and HEV-like viral isolates ^3^. HEV infection was originally thought to be solely restricted to humans, causing a self-limiting hepatitis ^4,5^. However, the discovery of HEV in swine in 1997 suggests that HEV has a much wider host range and is actually zoonotic ^6^. Currently, hepatitis E cases are frequently reported in developed countries and exhibit expanding host ranges ^3^. Moreover, chronic HEV infection, HEV-related acute hepatic failure, and extrahepatic manifestations caused by HEV have been frequently reported in recent years ^7–10^, suggesting a complicated mechanism underlying HEV-related diseases, especially for zoonotic HEVs.

HEV contains a 7.2 kb mRNA-like genome, which is capped and polyadenylated^11^. To date, three well-recognized open reading frames (ORFs) have been identified in all HEV isolates regardless of the genotypes or species ^12,13^, while the presence of an additional ORF4 has only been reported in *HEV-1*^14^ The HEV-ORF1 protein is translated directly from the viral genome, and it is a non-structural polyprotein that acts as the viral replicase ^15^. ORF2 and ORF3 which are partially or completely overlapped need to be transcribed into subgenomic RNAs for the protein translation ^16^. HEV-ORF2 encodes the major capsid protein, whereas HEV-ORF3 encodes a multifunctional protein that is probably a class I viroporin essential for virion release ^17^ The additional ORF4 in *HEV-1* is embedded within ORF1, and its protein production is driven by an internal ribosome entry site (IRES) initiation mechanism^14^. Existing evidence suggested that ORF4 protein stimulates ER stress to promote HEV replication and interacts with multiple ORF1 domains to form a complex enhancing the viral RdRp activity ^14^.

HEV-ORF3 encodes a 114-aa protein with a predicted molecular weight of 13 kDa (vp13) ^16,18^. Basic sequence analysis of HEV-ORF3 protein (Sar55 strain) reveals two hydrophobic domains within its N-terminal half and two proline-rich domains within its C-terminal portion ^19,20^, along with a MAP kinase phosphorylation site (Ser71) in the first proline-rich domain ^21^. Although the ORF3 protein is not required for viral RNA replication *in vitro* ^22^, it is essential for HEV virion release from infected cells and indispensable for HEV infection *in vivo*^18,23,24^. Studies have shown that a PSAP motif is required for the formation of membrane-associated HEV particles, a process that relies on the association of ORF3 protein with lipids ^25,26^. HEV-ORF3 forms an ion channel that shares key structural features with class I viroporins, and its ion channel function is required for virus particle release from cells during infection ^27^. This observation aligns with the reported putative role of pORF3 ^24^ and is consistent with the evidence that HEV-ORF3 interacts with tumor susceptibility gene 101 (TSG101), a component of the endosomal sorting complex involved in the generation of the endosomal sorting complex required for transport (ESCRT)^26^. The ESCRT is involved in budding of enveloped viruses and its formation leads to the biogenesis of quasi-enveloped HEV particles ^28–31^. Besides playing essential roles in virion release, studies have demonstrated that the HEV-ORF3 protein also interferes with multiple host cell signaling pathways, including pathways involved in host innate immunity ^32–34^,

In our previous investigations on HEV-ORF3, we observed that ORF3 from HEV-1 Sar55 strain overexpressed in HEK293T cells was able to form high-molecular-weight aggregates (unpublished data) consistent with a later observation that the ORF3 protein from HEV-3 KernowC1-p6 strain forming SDS-resistant polymers ^27^. The high-molecular-weight aggregates of HEV-ORF3 are also SDS-resistant, and this characteristic mirrors the property of LEF10, the first virus-encoded prion protein identified from a DNA virus belonging to the family *Baculoviridae* ^35^. Prions are self-propagating protein conformers that were originally discovered in association with neurodegenerative diseases in mammals. In recent years, it has been emerged that prions are not exclusive to mammalian species nor necessarily pathogenic. Prion-forming proteins are widely spread in plant, cellular microbes and acellular microorganisms, and are structurally and functionally diverse. In addition to their pathogenic roles, they act as epigenetic information carriers that have important regulatory functions ^36–39^.

In this study, by employing a well-characterized yeast reporter system, we demonstrate that the prion domain of yeast prion protein Sup35 can be functionally replaced by HEV-ORF3, supporting the hypothesis that the HEV-ORF3 protein behaves as a virus-encoded prion. Furthermore, we identify a candidate prion-forming domain in ORF3 and generate an ORF3 mutant with reduced aggregation propensity. The role of the prion property of ORF3 in HEV infection was also investigated *in vitro* in cultured cells and *in vivo* in an animal model.

## 2. Material and Methods

### 2.1 Cells and viruses

S10-3 cells, a subclone of Huh-7 hepatoma cell line ^16^, HepG2/C3A cell, HEK-293T cells and HeLa cells were maintained in Dulbecco’s Modified Eagle Medium (DMEM; Thermo Fisher Scientific, Waltham, MA, United States) supplemented with 10% FBS (Thermo Fisher Scientific). Full-length RNA of HEV-3 KernowC1 p6 strain (M74506.1) was obtained by *in vitro* transcription from plasmid pSK-HEV-kernowC1 p6 digested with *MluI* ^40^, using AmpliCap-Max T7 High Yield Message Maker Kit (Cellscript, Madison, WI, USA). The capped RNA yielded from i*n vitro* transcription was further purified using TRIzol^™^ Reagent (Thermo Fisher Scientific) according to the manufacturer’s protocol.

Transfection of DNA plasmids into different mammalian cells was conducted using FuGENE^®^ HD (Promega, Madison, WI, USA) according to the manufacturer’s instructions. Transfection of HEV-RNA was conducted using an optimized protocol with DMRIE-C (Thermo Fisher Scientific). Briefly, after S10-3 cells reaching 70% confluence in a 12-well plate, the cell culture medium was discarded and the monolayer cells were washed with PBS for twice, followed by addition of 0.5 mL serum-free Opti-MEM (Thermo Fisher Scientific) to each well. For each well, 1 μg RNA was added into 50 μL Opti-MEM and mixed evenly with 4 μL DMRIE-C. After incubation at room temperature for 20 minutes, the mixture was added to cells and 1 mL DMEM with 10% FBS was added to each well at 5 hours post transfection (hpt). HEV-RNA transfected cells were cultured at 37^□^C for 7 days. HEV-3 KernowC1 p6 virus was rescued from the transfected cell culture supernatant and concentrated using a previously described protocol ^41^, and the virus was used to inoculate HepG2/C3A cells to generate the cells stably infected with HEV.

For shuffling studies and phenotypic testing in yeast, *Saccharomyces cerevisiae* strain LJ14 (*MATa ade1-14 trp1-289 his3D-200 ura3-52 leu2-3,112 SUP35::loxP p[SUP35-URA3][PSI+]*) was utilized. Yeast cells were cultured in yeast extract peptone dextrose (YPD) and synthetic dropout (SD) Media at 30°C.

### 2.2 Ethics statement and animal studies

Specific-pathogen-free (SPF) male Mongolian gerbils (*Meriones unguiculatus*) at 8–10□weeks of age were purchased from the Experimental Animal Center at the Zhejiang Academy of Medical Sciences (Hangzhou, China). Prior to inoculation, all animals were confirmed to be seronegative for HEV by a commercial ELISA kit (Wantai Biological Pharmacy Co., Beijing, China). The animal protocol was reviewed and approved by the Animal Welfare Committee of Northwest A&F University. All gerbils were monitored on a daily basis for any clinical signs. All effort was made to minimize the suffering of animals and euthanasia was performed if the humane end-point was reached according our protocol. Purified HEV virions (WT and ORF3-F10S) obtained from cell culture media of HEV-3 KernowC1-p6 WT or ORF3-F10S stably infected HepG2/C3A cells were used to inoculate gerbils (n=5) at a dose of 1 × 10^9^ RNA copies of HEV in 1mL volume for each gerbil.

Gerbil serum samples and fecal samples were collected and tested for HEV-RNA at 7, 14, 21, 28, 35, and 42□ days post-infection (dpi) by real-time RT-PCR. Anti-HEV IgG levels in serum samples were monitored by ELISA using recombinant ORF2-p239 of HEV-3 KernowC1-p6 as the coating antigen. All the gerbils were sacrificed at 43dpi. Samples of liver, bile, and spleen were collected at necropsy and tested for the presence of HEV RNA by real-time RT-PCR. For histological examinations, additional 9 gerbils (n=3) were infected by HEV-3 KernowC1-p6 WT or ORF3-F10S viruses for two weeks and liver samples from virus infected animals and control animals were dissected, fixed in 4% paraformaldehyde for 12 h at 4°C, dehydrated in a graded series of ethanol, embedded in paraffin, and sliced into 6-μm-thick sections. The sections were subjected to histological examinations by hematoxylin and eosin (H&E) staining.

### 2.3 Plasmids and antibodies

The ORF3 cDNA sequences of HEV-1 Sar55 strain (GenBank Accession: AF444002.1) and HEV-3 KernowC1 p6 strain were amplified from infectious clones pSK-HEV2 and pSK-HEV-KernowC1 p6, followed by ligation to pCAGEN plasmids to fuse with an N-terminal MYC tag. The ORF3 sequence of HEV-2 Mexican strain (GenBank Accession: HQ709170.1) and HEV-4 Chinese strain (GenBank Accession: KF176351) were artificially synthesized and ligated to pCAGEN plasmids fused with an N-terminal MYC tag as well. The ORF3 of HEV-3 KernowC1 p6 strain (HEV-3 ORF3) was ligated to pET28a for recombinant protein expression in *E. coli*. Gene fragments for triple alanine screening and single amino acid (aa) mutation of HEV-3-ORF3 were artificially synthesized and ligated to pCAGEN plasmid. Generation of Bimolecular Fluorescence Complementation (BiFC) plasmids containing ORF3 sequence (HEV-3 KernowC1-p6 strain) were conducted by subcloning to BiFC vectors as previously described ^42^.

To investigate whether the ORF3 protein shows a prion-like aggregation trait in yeast cells, HEV ORF3 coding sequences were cloned into pUKC1620 between the *XbaI* and *Eco*RI sites to create pUKC1620-derived plasmids, such that the fused protein can be expressed under the control of the *SUP35* promoter. The plasmid p6431 (*CUP1*, GFP) was obtained from Professor Susan Lindquist at MIT. The HEV ORF3 fragments were cloned in-frame with the GFP coding sequence into plasmid p6431 between the *Sac*II and *NheI* sites, under the control of the yeast *CUP1* promoter. All the plasmids generated in this study were confirmed by sequencing.

Development of HEV-ORF2 specific mouse monoclonal antibody (Mab) was previously described ^43^. Recombinant HEV-ORF2-239 protein of HEV-1 Sar55 strain (p239) was prepared as previously described for rabbit polyclonal antibody production ^44^. The immunization of rabbits and affinity purification of p239-specific polyclonal antibody from immunized rabbits were conducted by Genscript Co. Ltd (Nanjing, China). The HEV-ORF3 rabbit polyclonal antibody were developed using an artificial peptide (aa sequence: CPSAPPLPHVVDLPQLGPRR) mapping to the C-terminal of HEV-1 Sar55 strain ORF3, which was conjugated to keyhole limpet hemocyanin (KLH). The peptides synthesis, KLH conjugation, rabbit immunization and antibody purification were conducted by Genscript Co. Ltd as well. The HEV-3 (KernowC1 p6) ORF3 specific Mab was developed by immunization of mice with recombinant HEV-3-ORF3 protein fusing with MBP tag followed by mice immunization, hybridoma fusion and screening. The expression of MBP fused HEV-3-ORF3 protein was conducted using a previously described method ^45^. Sup35C specific antibody was purchased from Santa Cruz Biotechnology (# SC-25915), and secondary antibodies used in this study were purchased from commercial suppliers.

### 2.4 Plasmid shuffling and phenotypic analysis in yeast

To produce a yeast strain with the ORF3-Sup35MC fusion protein as the only functional source of Sup35, the yeast LJ14 strain was transformed with ORF3-Sup35MC plasmid variants by standard PEG/LiAc/ssDNA transformation method ^46^. The transformed yeast cells were plated onto synthetic medium lacking uracil and histidine at grown at 30°C for 3 days, then transferred to YPD medium supplemented with 1 mg/ml 5-fluoroorotic acid (Sangon Biotech) to eliminate the *SUP35* maintainer plasmid p[*SUP35-URA3*].

To investigate if the premature UGA stop codon in the *ade1-14* allele was read through, the colonies that had lost p[SUP35-URA3] were grown overnight in YPD medium at 30°C, and the diluted cultures were respectively streaked onto rich 1/4 YPD agar, rich 1/4 YPD agar containing 5 mM guanidine hydrochloride and SD-Ade plate. In cells with a [*PSI*^+^] phenotype, the Sup35 protein aggregates resulting in impaired translation termination ability, which in turn results in the readthrough of the *ade1-14* stop codon and the synthesis of full-length Ade1 protein (N-succinyl-5-aminoimidazole-4-carboxamide ribotide synthetase). Such readthrough results in white colonies forming on both rich 1/4 YPD agar and SD-Ade agar plates. In contrast, in cells with a [*psi*^-^] phenotype, the *ade1-14* premature stop codon is not read through and therefore the cells form red colonies on rich 1/4 YPD agar and are not able to grow on SD-Ade medium.

To knock out the *HSP104* gene from LJ14 genome by homologous recombination, the endogenous *URA3* gene and its promoter fragments were amplified from plasmid p6431 by PCR, with the flanking sequences homologous to *HSP104* sequences added to both arms of the PCR fragment. Then, the linear replacement fragment was transformed into the yeast cells expressing full-length Sup35 or ORF3-Sup35MC by standard PEG/LiAc/ssDNA transformation. The transformed cells were cultured on SD-URA medium at 30°C for 3 days and then streaked on 1/4 YPD media for phenotypic observation.

### 2.5 Immunoblotting analysis

Cells were harvested by Laemmli sample buffer (Bio-Rad Laboratories, Hercules, CA, USA) containing 2% sodium dodecyl sulfate (SDS) and separated by SDS-polyacrylamide gel electrophoresis (SDS-PAGE) as previously described ^47^. For Semi-Denaturating Detergent Agarose Gel Electrophoresis (SDD-AGE), cells were harvested using lysis buffer (100 mM Tris 7.5, 50 mM NaCl, 10 mM BME) supplemented with protease inhibitors cocktail (Sigma-Aldrich, St. Louis, MO, USA) and separated by 2% agarose gel. Separated proteins were transferred onto PVDF membranes, probed with antibodies against different targets followed by HRP-conjugated anti-mouse or anti-rabbit IgG secondary antibodies (Thermo Fisher Scientific), and then the protein bands were visualized using enhanced chemiluminescent (ECL) substrate (Bio-Rad Laboratories). Chemiluminescence signal acquisition was conducted using a ChemiDoc MP Imaging System (Bio-Rad Laboratories) and the data analysis was performed using Image Lab software (Version 5.1, Bio-Rad Laboratories).

### 2.6 Fluorescence-activated cell sorting (FACS)

To analyze the fluorescence of BiFC assay, HEK-293T cells were transfected with BiFC plasmids for 24 hours, and then treated with trypsin to make single cell suspension. Flow cytometry assay was performed by using a FACSAria^™^ III cell sorter (BD Biosciences, San Jose, CA, USA) and the FACS data were analyzed using FlowJo software, version 10.0.7 (Tree Star, Ashland, Oregon, USA).

### 2.7 Immunofluorescence assay (IFA) and cell imaging

S10-3 cells were fixed at 7 days post transfection of different HEV-RNAs by using 2% paraformaldehyde for 15 min at RT, and then permeabilized with PBS containing 0.5% Triton X-100 (Sigma-Aldrich, St. Louis, MO, United States). Next, cells were probed with rabbit anti-p239 polyclonal antibodies and Alexa Fluor^®^555-labeled goat anti-rabbit IgG (Thermo Fisher Scientific). Cellular nuclei were counterstained with 4□, 6-diamidino-2-phenylindole (DAPI; Thermo Fisher Scientific) at 37 °C for 10 min. Images were captured under a Leica DM1000 fluorescence microscope (Leica Microsystems, Wetzlar, Germany) and processed using Leica Application Suite X (Version 1.0, Leica Microsystems).

To visualize ORF3-GFP proteins in yeast cells, the p6431-ORF3s plasmid was transformed into yeast strain LJ14-SUP1 by standard PEG/LiAc/ssDNA transformation method and cultured in SD-URA medium to logarithmic growth phase. CuSO_4_ was added to a final concentration of 25 μM to induce the expression of GFP fusion protein. Yeast cells were imaged at 4 h post-induction. Confocal images were cropped and processed with the ImageJ Software.

### 2.8 RNA isolation and quantitative real-time PCR (qPCR)

Total RNA was extracted from cells, serum or fecal samples using TRIzol^™^ Reagent (Thermo Fisher Scientific) in accordance with the manufacturer’s instructions. Reverse transcription were conducted using a PrimeScript RT reagent Kit (TaKaRa, Dalian, China). HEV RNA from different samples was detected using 2 × RealStar Power SYBR Mixture (Genstar, Beijing, China) or PerfectStart II Probe qPCR SuperMix following the manufacturer’s instructions. Transcripts for tubulin were amplified from the same cDNA to normalize total RNA input. For absolute quantification of HEV-RNA copies, the infectious clone plasmid for HEV-3 KernowC1-p6 was used for standard curve generation. Primers and Taqman probe used for qPCR and corresponding DNA sequences are listed in supplementary Table 1.

### 2.9 The enzyme-linked immunosorbent assay (ELISA)

For the evaluation of secreted HEV-ORF2 from HEV-3 KernowC1-p6 infected cells, 96-well Polystyrene Microplates (Corning Inc., Corning, NY, USA) were coated overnight with 1 μg/well of HEV-3 ORF2 specific Mab-2G8 in 100 μL of ELISA coating buffer (carbonate buffer, pH = 9.5-9.6) at 4 °C. Unbound Mab was removed by washing wells with PBS-T buffer and wells were further blocked by addition of PBS-T buffer containing 5% skim milk for 3 hours at 37 °C. Next, 100 μL cell culture supernatant was added and followed by incubation for 1 hour at 37 °C. After washing with PBS-T buffer for three times, 100 μL of PBS-diluted rabbit anti-ORF2 polyclonal antibody (1μg/mL) was added and incubated for 1 h at 37 °C. The proteins associated with the primary antibodies were further incubated with HRP-conjugated goat anti-rabbit secondary antibodies (Transgene) (1:5,000 dilution in PBS) for 1 h at 37 °C. Results were visualized after addition of 3,3’,5,5’-tetramethylbenzidine (TMB) substrate (Tiangen Biotech, Beijing, China). A standard curve for the HEV-ORF2 sandwich ELISA was generated using a serial dilution of recombinant ORF2-p239 protein of HEV-KernowC1-p6 strain (ORF2-p239) (2000, 1000, 500, 333, 111, 55.5, 27.75, 13.875, 0 pg/mL), and the figure was plotted using GraphPad Prism version 5.0 (GraphPad Software, San Diego, CA, USA). Absorbance values were measured using a VictorX5^™^ Multilabel Plate Reader (Perkin Elmer, Waltham, MA, USA) at a wavelength of 450 nm.

For the evaluation of serum antibody against HEV-ORF2, the 96-well Polystyrene Microplates (Corning Inc.) were coated with 400 ng/well ORF2-p239 in 100 μL of ELISA coating buffer (carbonate buffer, pH = 9.5-9.6). Serum samples from Mongolian gerbil were diluted 20-fold in PBS-T and added into the coated plate to incubate with the antigen for 1 hour at 37 °C. HRP-conjugated goat anti-Mongolian gerbil antiserum (Solarbio Life Science, Beijing, China) (1:5,000 dilution in PBS) was used as the secondary antibody.

### 2.10 Statistical analysis

Results were analyzed using GraphPad Prism version 5.0 (GraphPad Software, San Diego, CA, USA). Statistical significance was determined using either the Student’s t-test for comparison of two groups or by one-way analysis of variance (ANOVA) for more than two groups. A two-tailed *P* value < 0.05 was considered statistically significant.

## 3. Result

### 3.1 The HEV-ORF3 protein forms high molecular weight aggregates in mammalian cells

The HEV-ORF3 protein is a multifunctional protein that plays an indispensable role during HEV infection. In our previous studies on HEV-ORF3 from the HEV-1 Sar55 strain, we observed that the protein, when transiently overexpressed in different cell lines (HEK-293, S10-3, HeLa cells), formed SDS-resistant high-molecular-weight aggregates (unpublished data). However, due to the lack of an effective cell culture model and ORF3-specific monoclonal antibody (Mab), it was unknown if the SDS-resistant ORF3 oligomers existed in HEV infected cells during its natural replication cycle or such aggregation was simply an artifact of protein overexpression. Therefore, following the establishment of a cell-adapted HEV-3 KernowC1-p6 *in vitro* culture system and the generation of an HEV-3 KerowC1-p6 specific ORF3 Mab, we were able to examine the ORF3 protein natively expressed in HEV-3 KernowC1 p6 infected cells. As shown in Figure 1A, after the transfection of S10-3 cells with *in vitro* transcribed HEV-RNA, SDS-resistant polymers of the ORF3 protein could be detected. By semi-denaturing detergent agarose gel (SDD-AGE) electrophoresis, a common assay for the detection of amyloid-forming proteins^48^, ORF3 was detected as a diffuse band, confirming the formation of SDS-resistant polymers. In HepG2/C3A cells which were stably infected with HEV, similar aggregates of ORF3 protein were detected by both SDS-PAGE and SDD-AGE (Figure 1B).

**Figure 1.**
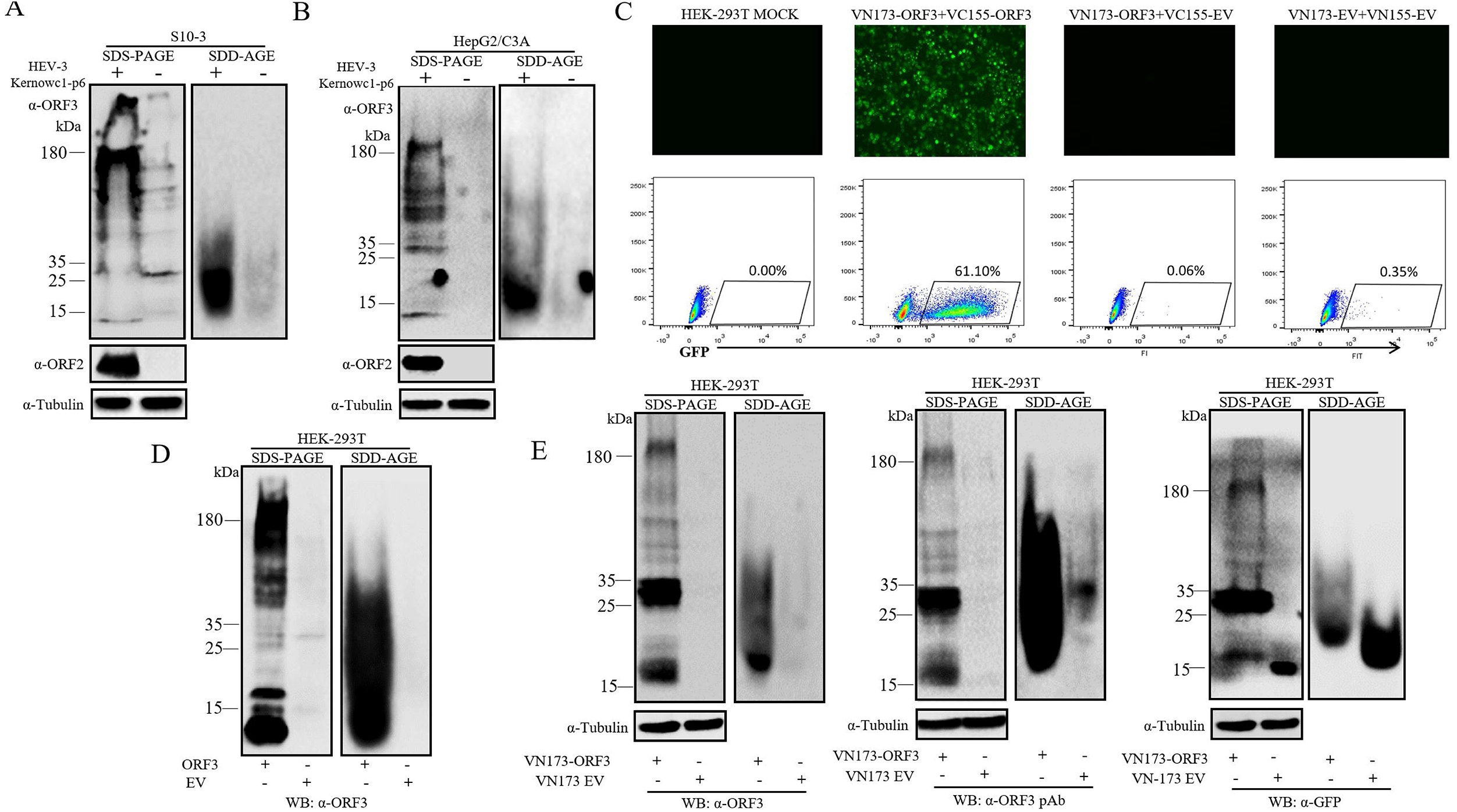
The HEV-ORF3 protein forms SDS-resistant high-molecular-weight aggregates in mammalian cells. **A.** SDS-PAGE and SDD-AGE analyses of HEV-ORF3 in S10-3 cells transfected with *in vitro* transcribed HEV-3 KernowC1-p6 RNA. The cells were harvested at 7 days post transfection using Laemmeli sample buffer for SDS-PAGE or the lysis buffer for SDD-AGE. After the separated proteins were transferred to PVDF membrane, ORF3 were detected with an HEV-3 ORF3 specific-monoclonal antibody (Mab, α-ORF3). HEV-ORF2 in the cell lysates was detected by rabbit anti-HEV-ORF2 polyclonal antibody α-ORF2) as infection control, and tubulin was probed as protein loading control. **B.** SDS-PAGE and SDD-AGE analyses of HEV-ORF3 in infected HepG2C3A cells. **C.** Detection of ORF3 self-interaction by BiFC assays. HEK-293T cells were co-transfected with the indicated plasmids expressing N-terminal Venus protein (VN173 EV), C-terminal Venus (VC155 EV), and HEV-ORF3 proteins fused with the tags (VN173-ORF3, VC155-ORF3). The fluorescence signals in the transfected cells were observed under fluorescence microscope and also examined by flow cytometry. **D.** SDS-PAGE and SDD-AGE analyses of overexpressed HEV-ORF3 in transfected HEK-293T cells **E.** SDS-PAGE and SDD-AGE analyses of overexpressed HEV-ORF3 tagged with VN173. Anti-HEV-3-ORF3 Mab (α-ORF3) and rabbit anti-HEV-3 polyclonal antibody (α-ORF3 pAb) were used for the detection of HEV-ORF3, and the fused tag was examined using anti-GFP polyclonal antibody (α-GFP).

To confirm that the aggregation of the ORF3 protein was a consequence of self-assembly rather than covalently attaching to other cellular proteins like ubiquitin, a bimolecular fluorescence complementation (BiFC) assay was conducted in HEK-293T cells. Strong florescence was observed only in the cells expressing the ORF3 fused to both the N-terminal and C-terminal Venus protein (Figure 1C), demonstrating the self-interaction of the HEV-ORF3 proteins. Consisted with the BiFC result, high-molecular-weight aggregation of ORF3 protein could be observed by both SDS-PAGE and SDD-AGE in HEK-293 cells overexpressing ORF3 protein without a tag (Figure 1D) or with the VN173 tag (Figure 1E, left panel). Immunoblotting the proteins separated by SDS-PAGE and SDD-AGE using an anti-ORF3 polyclonal antibody (Figure 1E, middle panel) and an anti-GFP tag polyclonal antibody (Figure 1E, right panel) confirmed the existence of SDS-resistant ORF3 polymers, although it seemed that the VN173 tag reduced the level of aggregation of ORF3 proteins. Taken together, these observations demonstrate that HEV ORF3 protein is able to form SDS-resistant high-molecular weight aggregates during infection in susceptible hepatoma cell lines as well as when the protein was overexpressed in HEK-293T cells.

### 3.2 HEV-ORF3 protein behaves as a prion in a yeast prion reporter system

Formation of SDS-resistant aggregates is a typical characteristic of prion and prion-like proteins ^48^. To verify the prion traits of HEV-ORF3 protein, a yeast prion reporter system based on the well-characterized prion phenotypes of the *Saccharomyces cerevisiae* translation termination factor Sup35 was utilized. In this system, when the N-terminal prion-forming domain (PrD) of Sup35 is functionally replaced by other PrD bearing proteins, it will endow yeast cells with a [*PSI*^+^]-like phenotype ^49^. Here, the PrD of Sup35 was substituted by HEV-3 ORF3 to generate a recombinant ORF3-Sup35MC fusion protein. Like the [*PSI*^+^] phenotype caused by the prion form of wild-type Sup35, the yeast strain harboring the recombinant ORF3-Sup35MC protein exhibited both [*ORF3*^+^] and [*orf3*^-^] phenotypes. In the yeast cells where Sup35 or ORF3-Sup35MC was soluble, protein synthesis is efficiently terminated at the premature UGA stop codon in the yeast *ade1-14* allele, which results in red colonies on the 1/4 YPD plate and a failure to grow on SD-Ade medium. By contrast, when the cells carry prion-like aggregates of these proteins they give rise to white colonies on both the 1/4 YPD plate and SD-Ade medium due to the read-through of the *ade1-14* premature stop codon (Figure. 2A and 2E). By counting the red and white colonies in three 1/4 YPD plates, 89.98% of the yeast cells expressing ORF3-Sup35MC formed [*orf3*^-^] colonies, and 10.02% formed [*ORF3*^+^] colonies.

**Figure 2.**
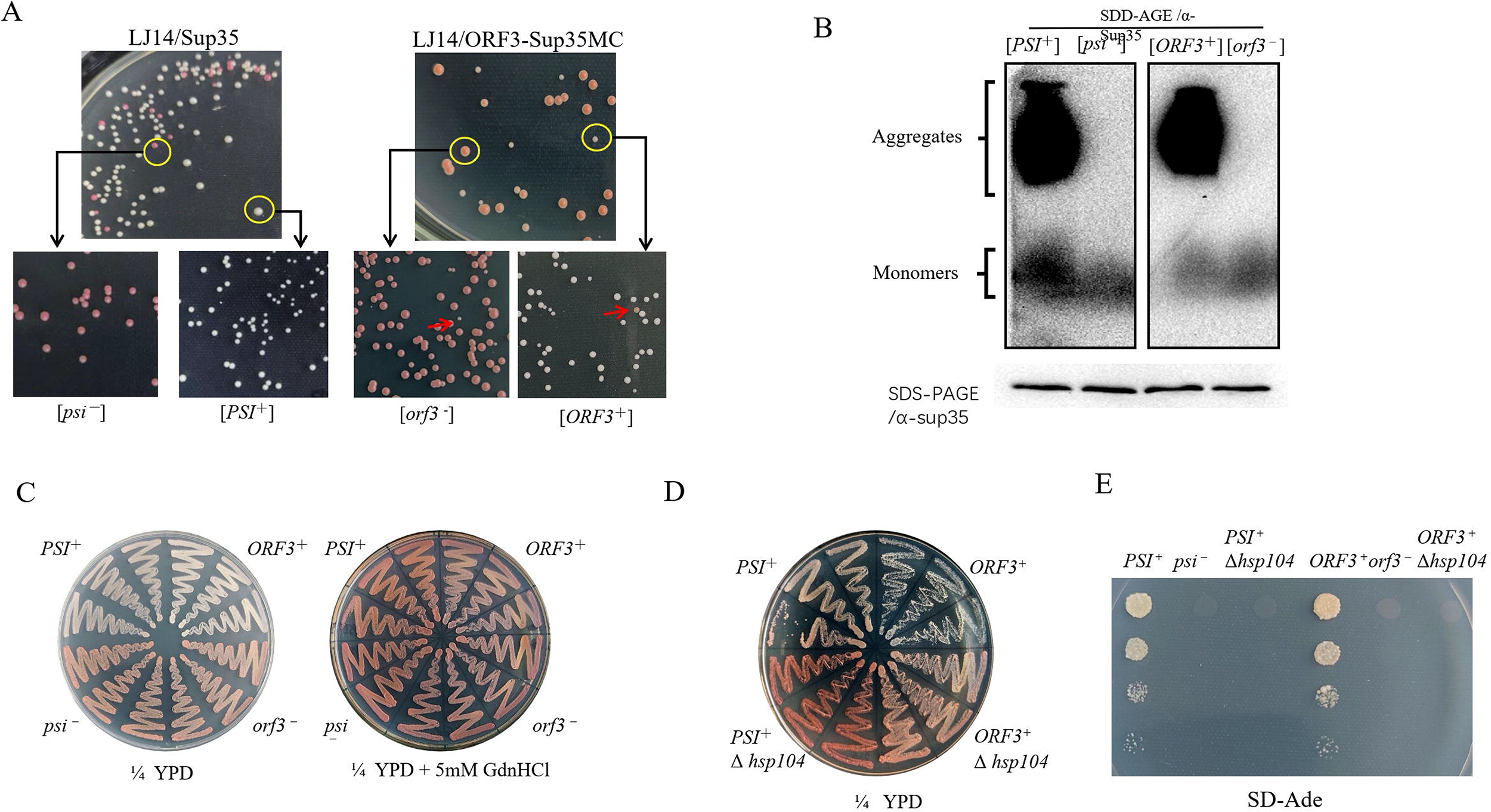
The HEV-ORF3 protein can functionally replace the prion-forming domain of Sup35 in a yeast prion reporter assay. **A.** HEV-ORF3-Sup35MC confers inheritable [*ORF3*^+^] and [*orf3*^-^] phenotypes to yeast cells. Yeast *ade1-14* cells expressing HEV-ORF3-Sup35MC were spread on complete (1/4 YPD) medium. [*ORF3*^+^] strains formed white colonies, which were distinguishable from [*orf3*^-^] strains having red pigment accumulated through the adenine biosynthetic pathway. Both phenotypes were stably inhertited and the switch of [*ORF3*^+^] and [*orf3*^-^] phenotypes occurred at a low frequency during propagation (indicated by red arrows). Normal yeast *ade1-14* cells expressing wild-type Sup35 were included as controls to show the typical [*PSI*^+^] and [*psi*^-^] phenotypes. **B.** Detection of SDS-resistant aggregates in [*ORF3*^+^] strains by SDD-AGE. The expression of HEV-ORF3-Sup35MC and Sup35 were probed by anti-Sup35 antibody (α-Sup35). The expression levels of HEV-ORF3-Sup35MC and Sup35 were examined by SDS-PAGE. **C.** The [*ORF3*^+^] phenotype on 1/4 YPD medium is cured by the treatment with guanidine hydrochloride (GdnHCl). **D.** The [*ORF3*^+^] phenotype on 1/4 YPD medium is curable by *hsp104* gene knockout. **E.** The [*ORF3*^+^] phenotype on medium lacking adenine (SD-Ade) is curable by *HSP104* gene knockout.

During the propagation of [*ORF3*^+^] and [*orf3*^-^] colonies, the white or red phenotype was stable through multiple cell generations although the switch of phenotype (0.89% from white to red, 1.81% from red to white) could be observed at a very low frequency (Figure. 2A). As the ORF3-Sup35MC fusion protein was expressed under the control of the *SUP35* gene promoter, this result indicated that the prion conformation of HEV-3 ORF3 was self-perpetuating and the low expression level was sufficient to maintain its prion state. Analysis by SDD-AGE confirmed that the ORF3-Sup35MC protein formed high-molecular-weight, SDS-resistant aggregates in [*ORF3*^+^] cells, whereas only monomers of this protein could be detected in [*orf3*^-^] colonies (Figure. 2B) consistent with the *in vivo* [*ORF3*^+^] and [*orf3*^-^] phenotypes.

Hsp104 is a molecular chaperone required for the heritability of [*PSI*^+^] and other yest prions ^50^. To test if Hsp104 is also essential for the maintenance of the [*ORF3*^+^] phenotype, an Hsp104-specific inhibitor guanidine hydrochloride (GdnHCl) ^51^ was applied to inactivate the chaperone function of Hsp104. By growing [*ORF3*^+^] yeast cells on rich medium containing 5□mM GdnHCl, the [*ORF3*^+^] phenotype was lost resulting in [*orf3*^-^] cells as is observed with [*PSI*^+^] cells (Figure. 2C). Further data on the deletion of the *HSP104* gene showed that the lack of Hsp104 eliminated the [*ORF3*^+^] phenotypes as confirmed by growth on both 1/4 YPD and SD-Ade medium (Figure. 2D and 2E), indicating that the [*ORF3*^+^] phenotype requires Hsp104 for the *de novo* induction and propagation of the phenotype. Such Hsp104-dependent characteristics have been observed for almost all identified yeast prions ^50,52^, which further demonstrates that the [*ORF3*^+^] phenotype was not caused by non-epigenetic effects but by the *bona fide* prion property of the ORF3 protein in yeast cells.

### 3.3 The ORF3 proteins from HEV-1 to HEV-4 have prion characteristics

Sequence analysis reveals that ORF3 proteins from different HEV genotypes within *Orthohepevirus A* are not completely conserved, especially in the C-terminal half of the protein ^53^, thus it is interesting to know if all HEV-ORF3 proteins from HEV-1 to HEV-4 possess prion characteristics. To answer this question, the *ORF3* genes from the four HEV genotypes were each cloned into the mammalian expression vector pCAGEN. The proteins were fused with a MYC tag so that they could be detected with both a MYC tag antibody and an anti-ORF3-peptide polyclonal antibody. As shown in Figure 3A, although the anti-ORF3-peptide polyclonal antibody did not react well with the ORF3 from HEV-2 (GT-2), all the ORF3 proteins overexpressed in HEK-293T cells were found to form high-molecular-weight, SDS-resistant aggregates as detected by SDD-AGE when probed with a MYC-specific Mab.

**Figure 3.**
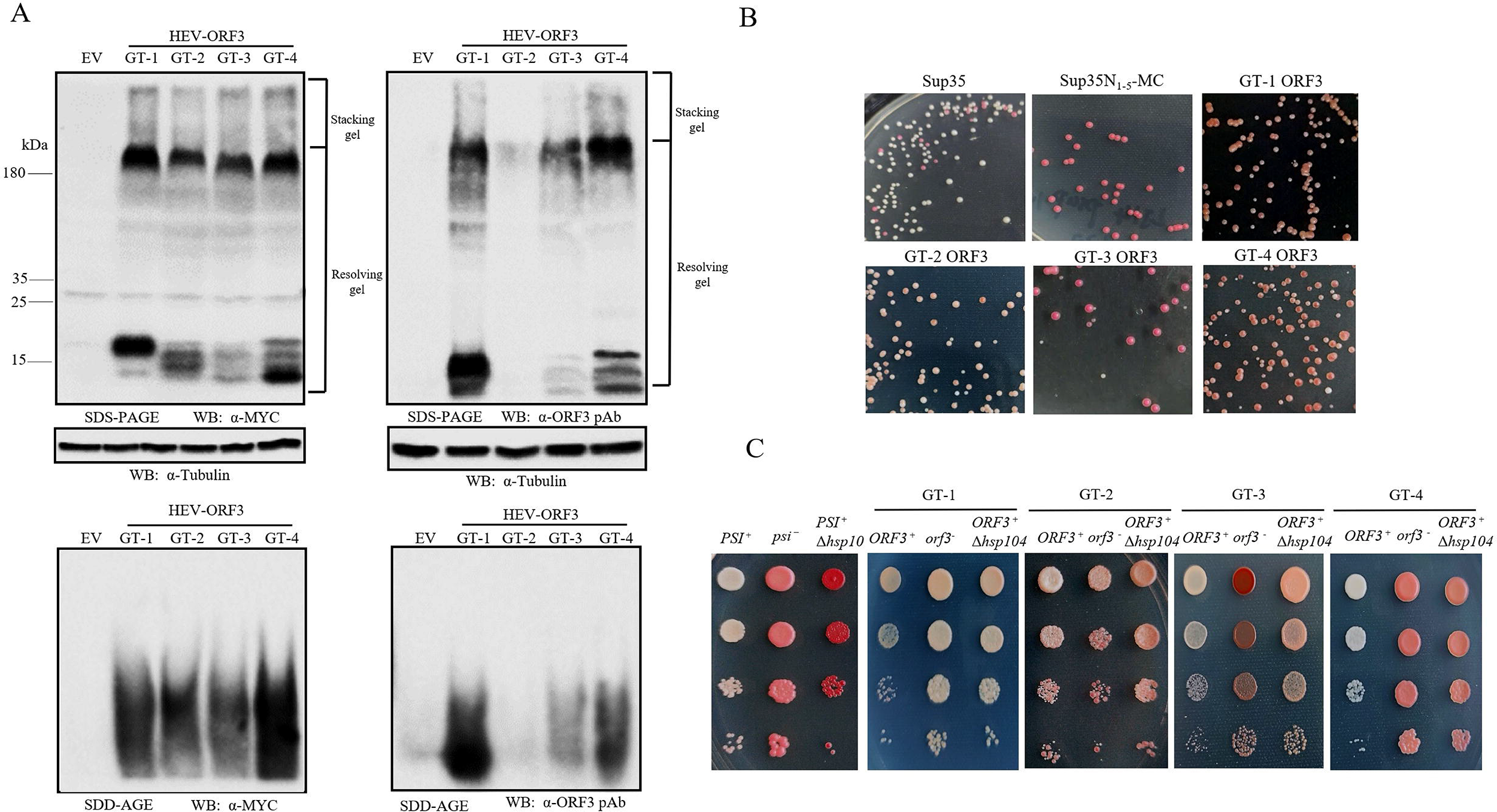
ORF3 proteins from different HEV genotypes display prion-like properties. **A.** ORF3 proteins form SDS-resistant polymers. HEK-293T cells were transfected with empty plasmid (EV) or plasmids encoding MYC-fused ORF3 proteins from 4 HEV genotypes (GT-1 to GT-4) and harvested at 24 hours post transfection for SDS-PAGE (upper panel) or SDD-AGE (lower panel). Western Blot analyses were performed using anti-MYC-Mab (α-MYC. left panel) and rabbit anti-HEV-ORF3 polyclonal antibody (α-MYC pAb. right panel). **B.** [*ORF3*^+^] and [*orf3*^-^] phenotypes endowed by HEV-ORF3-Sup35MC from 4 HEV genotypes (GT1 to GT4) on 1/4 YPD medium. **C.** Phenotypic curing of [*ORF3*^+^] phenotypes by the deletion of the *HSP104* gene. Serial dilutions of the indicated strains were spotted on the 1/4 YPD plate.

In yeast cells, all four ORF3 proteins could replace the function of the Sup35 PrD and generate [*ORF3*^+^] and [*orf3*^-^] phenotypes as shown for HEV-3 protein, although the two phenotypes for HEV-1 and HEV-2 were not as clearly defined as for HEV-3 and HEV-4 (Figure. 3B). By deletion of the *HSP104* gene, in all cases the [*ORF3*^+^] cells converted to the [*orf3*^-^] phenotype (Figure 3C), demonstrating that the [*ORF3*^+^] phenotype formed by ORF3 proteins of different HEV genotypes was Hsp104 dependent. Taken together, these data demonstrate that all ORF3 proteins from HEV-1 to HEV-4 have prion-like properties like the yeast prion Sup35 with the prion-like characteristic of ORF3 appearing to be conserved among the 4 genotypes of the HEV virus.

### 3.4 The N-terminus of HEV-ORF3 acts as prion-forming domain

Most prion proteins identified so far have glutamine/asparagine (Q/N)-rich, prion forming domains, but this feature is not present in the HEV ORF3 protein. Using bioinformatics tools to search for potential PrDs in ORF3 ^54–56^, we did not define a distinct candidate prion-forming domain (cPrD) by PLAAC (Supplementary Fig. S1A), but found several short amyloidogenic regions in the N-terminal part of ORF3 protein by AGGRESCAN and FoldAmyloid (Supplementary Fig. S1B and S1C). To identify the cPrD of HEV-ORF3, we replaced the Sup35 PrD with 5 different ORF3 truncations and found that the N-terminal 25 amino acids (N25) were enough to confer the [*ORF3*^+^] phenotype in yeast (Fig. 4A and Supplementary Fig. S1D). Therefore, N25 of HEV-ORF3 harboring the first potential amyloid formation region predicted by AGGRESCAN and FoldAmyloid algorithms was defined as the cPrD of HEV-ORF3.

**Figure 4.**
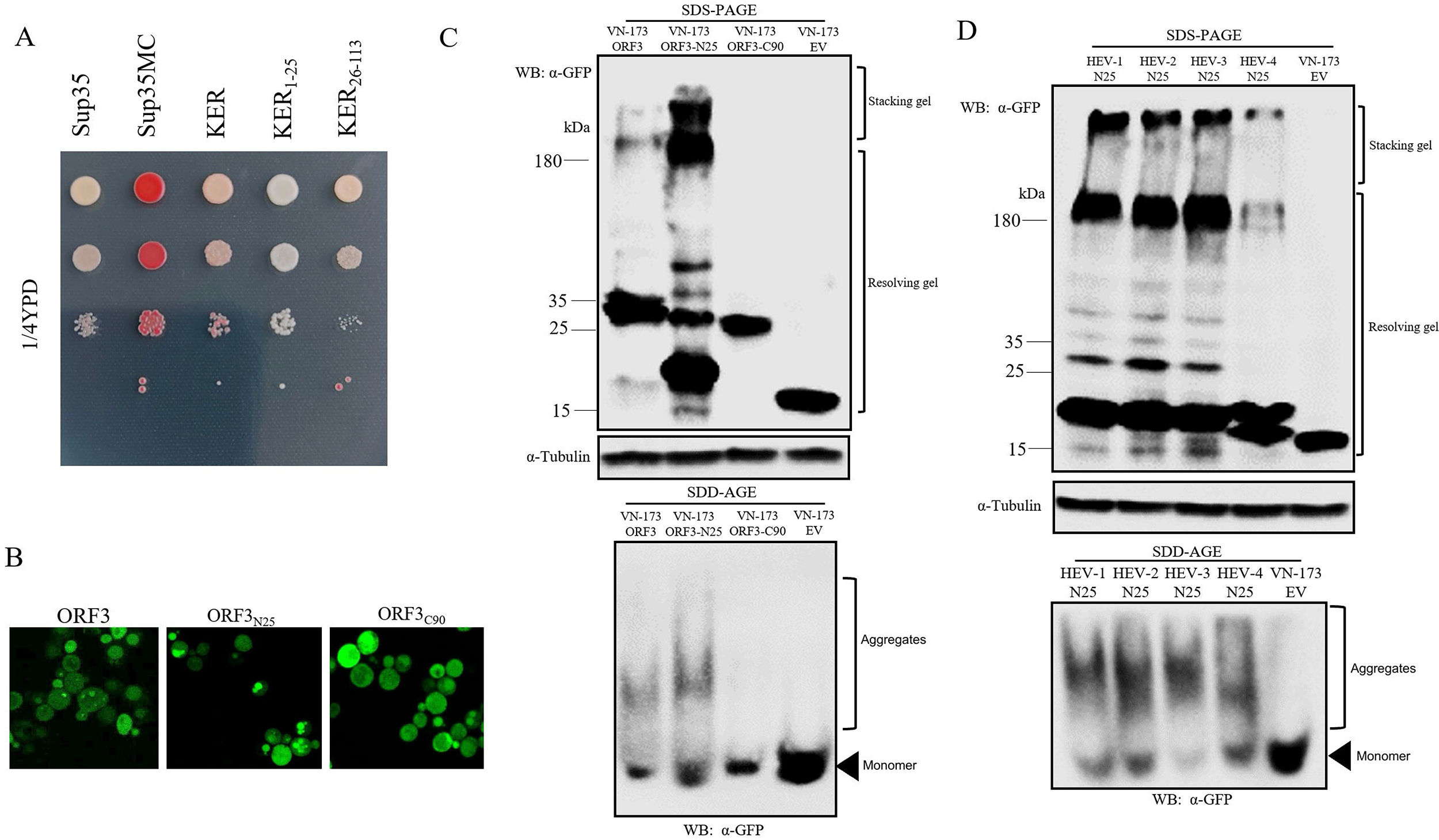
Identification of the N terminal 25 amino acids (N25) region as the candidate prion-forming domain (cPrD) of HEV-ORF3. **A.** N25-Sup35MC confers [*ORF3*^+^] phenotype to yeast cells on 1/4 YPD medium. **B.** ORF3N25-GFP produces big foci in yeast cells. In contrast. ORF3C90-GFP is evenly distributed. **C.** ORF3N25 fused with VN173 tag forms SDS-resistant polymers by SDS-PAGE and SDD-AGE analyses. HEK-293T cells were transfected with the indicated plasmids and the protein expression was examined at 24 hours post transfection. The ORF3 protein used in figure A-C is from HEV-3 KernowC1-p6. **D.** HEV-ORF3 proteins from 4 HEV genotypes (GT1 to GT4) form SDS-resistant polymers in HEK-293T cells by SDS-PAGE and SDD-AGE analyses. The protein was detected using an anti-GFP polyclonal antibody (α-GFP).

In a previous investigation on the interaction of ORF3 from HEV-1 Sar55 with the host cell microtubules, it was found that the ORF3 N-terminus was essential for the formation of microtubule-like filaments, but the fusion protein only containing the N-terminal ORF3 formed aggregated dots in the cells ^19^, which is very similar to the aggregated form of AcMNPV LEF10 protein ^35^. Here, in transformed yeast cells, we observed that ORF3_N25_-GFP produced larger punctate aggregates than ORF3-GFP while ORF3_C90_-GFP (containing the C-terminal 90aa of ORF3 protein) was evenly distributed throughout the cells (Figure 4B), consistent with the observation in human cells^19^.

To investigate if the N25 domain could form SDS-resistant aggregates, N25 and C90 were expressed as fusion proteins with a VN173 tag in HEK-293T cells. ORF3-N25 demonstrated stronger aggregation propensity than the full-length ORF3 protein, whereas ORF3-C90 protein lacking N25 did not form SDS-resistant high-molecular-weight aggregates (Figure 4C), confirming that N25 was the cPrD of HEV-ORF3. Further analysis of N25 from HEV-1, 2 and 4 showed that the N terminal potential amyloid formation region of these genotypes also formed SDS-resistant aggregates, like that of the N25 sequence originating from the HEV-3 KernowC1 strain. These data suggest that all ORF3 proteins from the four HEV-genotypes possessed similar cPrDs and prion-like characteristics.

Since prion-like protein has never previously been identified from RNA viruses, to understand the biological significance of the prion-forming character of HEV-ORF3, we tried to generate PrD-domain/function negative mutants. First, triple alanine screening for ORF3-N25 was conducted and 8 mutants (M1 to M8) with each bearing a set of alanine triplets were constructed. As shown in Figure 5A, these mutants demonstrated variable ability to form SDS-resistant aggregates, but M3 harboring mutations at positions 8-10 showed the lowest propensity to aggregate. (Figure 5A). In yeast cells, the M3-GFP mutant exhibited decreased aggregation tendency (Figure 5B) and expression of the ORF3(M3)-Sup35MC protein generated yeast cells with a very weak Ade^+^ phenotype closer to that observed for [*psi*^-^] cells on 1/4 YPD plate (Figure 5C).

**Figure 5.**
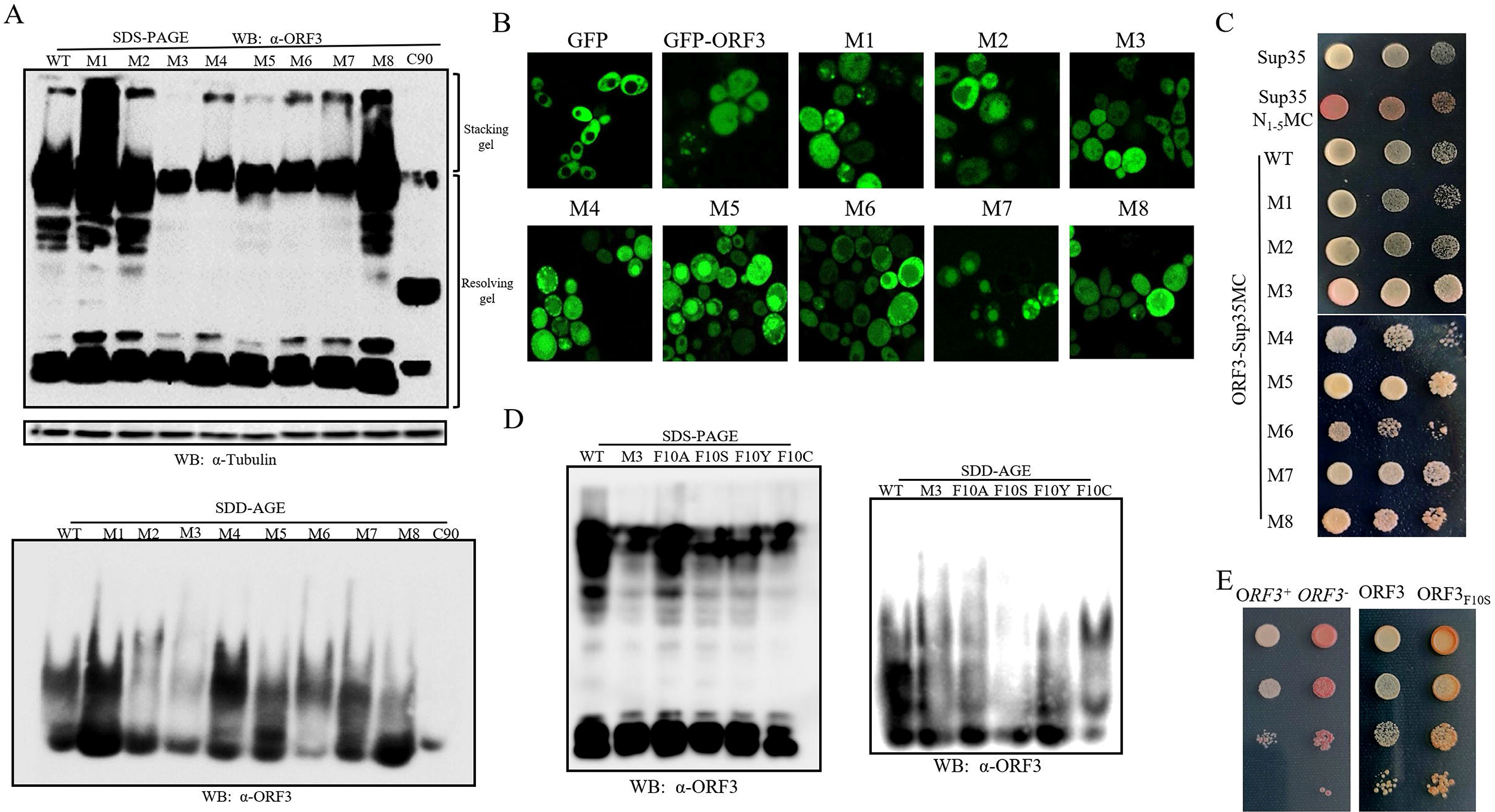
Phenylalanine at the amino acid position 10 (F10) is a key residue for the prion-forming trait of ORF3. **A.** SDS-PAGE and SDD-AGE analyses of ORF3 triple alanine mutants. HEK-293T cells were transfected with plasmids encoding wild type ORF3 (WT), 8 ORF3 mutants (M1 to M8) bearing triple alanine mutations from aa 2-25 of ORF3-N25, and VN173-fused C-terminal 90aa of ORF3 (C90). The cells were harvested at 24 hours post transfection for SDS-PAGE (upper panel) or SDD-AGE (low panel) analyses, and the truncated ORF3 proteins were probed using anti-ORF3 antibody (α-ORF3). **B.** Observation of the aggregation propensity of GFP tagged ORF3 triple alanine mutants in yeast cells by confocal microscopy. **C.** Phenotype analysis of yeast cells bearing ORF3-Sup35MC triple alanine mutants on 1/4 YPD medium. **D.** Identification of ORF3 F10S mutant with reduced ability to form SDS-resistant polymers. HEK-293T cells were transfected with plasmids encoding VN173-fused wild type ORF3 (WT) and the indicated mutants. The protein was detected using an anti-ORF3 Mab (α-ORF3). **E.** Phenotype analysis of yeast cells bearing ORF3F10S-Sup35MC on 1/4 YPD medium.

To further pin down the key amino acid residue involved in the prion aggregation of HEV-ORF3, single alanine residues were introduced at positions of aa 8-10 of ORF3, and the aggregation abilities of the mutants compared with the wild type HEV-ORF3 and M3 mutant. The ORF3_F10A_ produced the least amount of SDS-resistant aggregates among the three single aa mutants as defined by SDD-AGE and its aggregation tendency was close to that of the M3 mutant (Supplementary Fig. S2), suggesting that the phenylalanine at position 10 (F10) was a key amino acid involved in the prion-like behavior of the ORF3 protein. However, as HEV-ORF2 and ORF3 are partially overlapped, the F10A mutation in ORF3 will cause a nonsynonymous amino acid change in ORF2 when alanine is introduced into aa position 10 of HEV-ORF3 in KernowC1-p6 based infectious clones. Therefore, we tried to substitute the hydrophobic F10 with three different hydrophilic amino acid serine, tyrosine and cysteine as this would not introduce any amino acid change in ORF2. The F10S mutant showed dramatically reduced ability to form SDS-resistant aggregates when expressed in HEK-293T cells (Figure 5D). In the Sup35MC-based *in vivo* yeast assay, the F10S mutant had a phenotype closer to [*orf3*^-^] than was observed with the wild type ORF3 protein (Figure 5E), consistent with a reduced aggregation tendency.

### 3.5 Mutation of the ORF3 prion-forming domain prevents virus release from HEV-infected cells

To investigate the role of the ORF3 prion-forming function in the HEV replication cycle, a reverse genetic-based mutagenesis assay was carried out to generate an HEV infectious clone harboring the ORF3-F10S mutation. Similar to the wild type virus, the HEV-3 KernowC1-p6 ORF3-F10S virus was viable and could be successfully rescued from S10-3 cells after transfection of *in vitro* transcribed RNA (Figure 6A). Sequencing of the cDNA of virus recovered from cell culture supernatant confirmed the presence of the F10S mutation in ORF3 (Figure 6A). Using both SDS-PAGE and SDD-AGE, ORF3 aggregation in these HEV-RNA transfected cells was investigated, and a reduction of ORF3 aggregation was observed in the cells transfected with HEV-3 KernowC1-p6 ORF3-F10S (Figure 6B). Meanwhile, we also observed that the expression level of ORF2 protein in S10-3 cell transfected with KernowC1-p6 ORF3-F10S was obviously higher than the cells transfected with the wild type HEV RNA. To establish whether it was different viral RNA levels that accounted for the different ORF2 expression levels, the positive and negative strand viral RNAs were examined. In cells transfected with WT RNA and ORF3-F10S RNA, the negative strand HEV-RNA (template used for HEV-RNA synthesis) levels were similar in the two groups. Although the average level of the positive strand RNA in the F10S group was higher than the wild type group, there was no statistical significance between the two groups (Figure 6C). These data suggest that the F10S mutation did not affect RNA replication in the virus.

**Figure 6.**
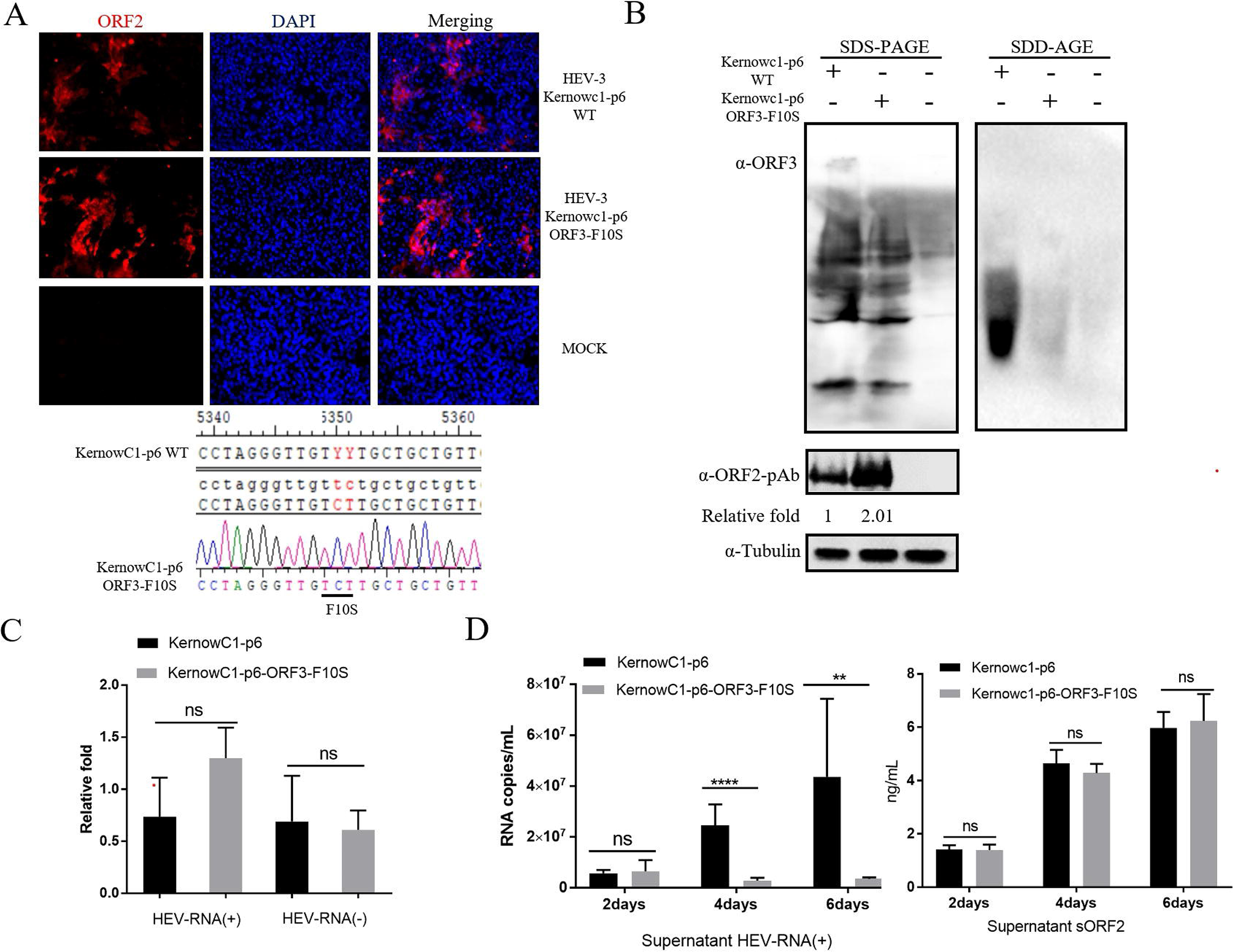
Prion-like aggregation of the HEV-ORF3 protein is involved in HEV virion release from infected cells. **A.** Rescue of HEV-3 KernowC1-p6 ORF3-F10S mutant virus by reverse genetic system. S10-3 cells were transfected with the indicated viral RNAs and incubated for 7 days. HEV replication was determined by immunofluorescence assay using anti-HEV-ORF2-specific Mab-2G8. Mock transfected S10-3 cells were included as negative controls. Cell culture supernatants were harvested for the amplification of full length ORF3 gene by RT-PCR. The mutation of F10S in ORF3 was confirmed by DNA sequencing. **B.** Examination of the SDS-resistant aggregates formed by ORF3 and ORF3-F10S in HEV-RNAs transfected S10-3 cells. The ORF3 proteins separated by SDS-PAGE (left panel) and SDD-AGE (right panel) were probed by HEV-3-ORF3 specific Mab (α-Mab). The expression of HEV capsid protein (ORF2) was detected using rabbit anti-HEV-ORF2-p239 polyclonal antibody (α-ORF2-pAb) to indicate the accumulation of virus particles. and tubulin was probed to normalize protein loading. **C.** Quantification of the levels of positive strand (HEV-RNA (+)) and negative strand (HEV-RNA (-)) HEV-RNAs in transfected S10-3 cells by qPCR. Experiments were repeated for at least three times. All data is presented as mean ± SD and subjected to Student’s *t*-test. “ns” means not significant. **D.** Quantification of the HEV-RNA (+) and secreted ORF2 protein levels in the supernatant of transfected S10-3 cells. Cell culture supernatant was harvested at 2, 4 and 6 days post RNA transfection. The plus strand viral RNA genome was determined by RT-qPCR. Secreted form of HEV-ORF2 was quantified by ELISA. Experiments were repeated for at least three times. All data is presented as mean ± SD and subjected to Student’s *t*-test. Significant differences between indicated groups are marked by “**” (p < 0.01) and “****” (p < 0.0001). whereas “ns” means not significant.

However, when the plus-stranded genomic RNA released into the cell culture supernatant was examined, a significant reduction in RNA level was observed in the ORF3-F10S mutant group at 4 and 6 days post transfection (Figure 6D), suggesting that the virion release process could be inhibited by the F10S mutation. It is possible that reduced virion release process caused the accumulation of assembled virion, which would account for a higher intracellular ORF2 protein level and an increased average level of plus strand viral RNA in ORF3-F10S mutant virus-infected cells. Meanwhile, since a recent report suggested that the majority of HEV-ORF2 could be directly secreted from infected HepG2/C3A cells before the assembly with genomic RNA ^57^, we performed a sandwich ELISA for the detection of the secreted form of the HEV-ORF2 protein in the cell culture supernatants rather than the “quasi-enveloped” particles. No significant difference was detected between the groups transfected with WT HEV-RNA and the ORF3-F10S mutant, suggesting that the F10S mutation did not influence the secretion of soluble ORF2, but probably only obstructed the release of assembled virus particles.

### 3.6 Mutation in the ORF3 prion-forming domain reduces microtubule stability in host cells

A conserved PSAP motif located at aa 95-98 of HEV-ORF3 is important for HEV-ORF3’s interaction with the tumor susceptibility gene 101 (TSG101) ^26^, a key component of the endosomal sorting complex required for transport (ESCRT), which is involved in budding and biogenesis of quasi-enveloped HEV particles ^28–31^. Using a Co-IP assay, we found that the ORF3-F10S mutant had a similar binding ability to TSG101 as the wild-type ORF3 (Figure 7A), indicating that the impaired virion release by F10S mutation was not a consequence of impaired interaction between TSG101 and ORF3-F10S.

**Figure 7.**
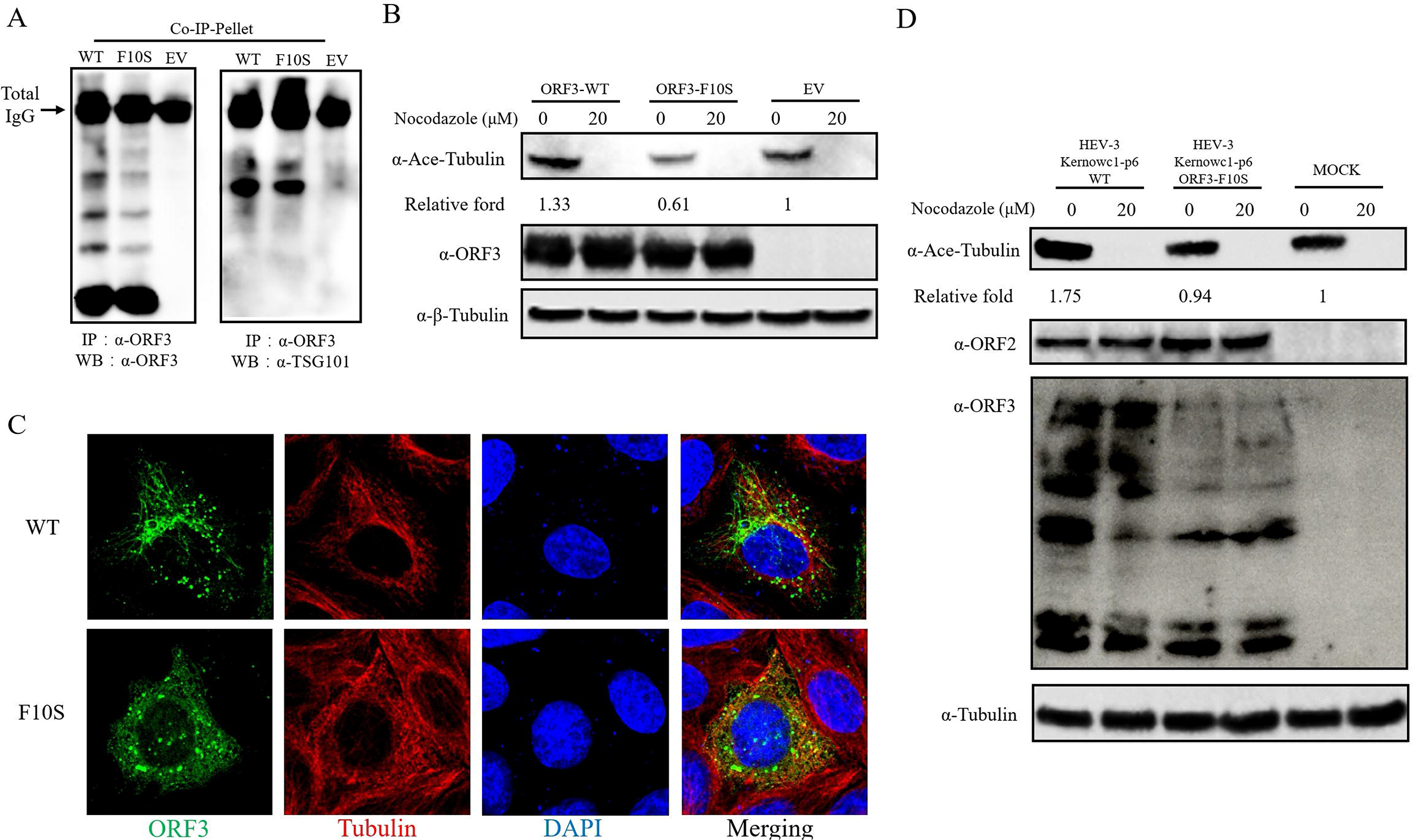
Mutation of HEV-ORF3 cPrD affects tubulin stability in HEV infected cells. **A.** F10S mutation does not interfere the interaction between TSG101 and ORF3. Transfected HEK-293T cells expressing the indicated ORF3 proteins were lysed in NP40 buffer for Co-IP assay using HEV-3-ORF3 specific Mab (α-ORF3). IP pellets pulled down by protein G agarose were harvest for Western blot analysis using α-ORF3 or anti-TSG101 Mab (α-TSG101). EV refers the negative control transfected with empty vector. **B.** Expression of ORF3-F10S mutant reduces the level of acetylated tubulin. HeLa cell were transfected with the indicated plasmids. Nocodazole treatment was carried out at 24 hours post transfection at a concentration of 20μM for 2 hours. Western blots were performed using anti-Ace-tubulin antibody and HEV-3-ORF3 specific Mab (α-ORF3). Total tubulin was probed to normalize the protein load. **C.** ORF3-F10S interferes with the formation of long filamentous structures by tubulin and ORF3. HeLa cells were transfected with Venus-tagged HEV-3-ORF3 (WT) and HEV-3-ORF3 bearing F10S mutant (F10S) for 24 hours. The expression of the fluorescent ORF3 fusion protein (Green channel) and tubulin probed by anti-tubulin antibodies (Red channel) were observed under a confocal microscope. **D.** F10S mutation abolishes the role of ORF3 in enhancing tubulin acetylation in virus infection. S10-3 cells were transfected with HEV-RNAs (KernowC1-p6 WT or KernowC1-p6 ORF3-F10S). Nocodazole treatment was carried out at 7 days post transfection at a concentration of 20μM for 2 hours. Western blots were performed using anti-Ace-tubulin antibody and HEV-3-ORF3 specific Mab (α-ORF3). Total tubulin was probed to normalize the protein load.

Tubulin acetylation is a well-established marker of microtubule stability ^58^. A previous report demonstrated that enhanced acetylation of tubulin in influenza A virus (I AV)-infected epithelial cells was correlated with increased virion release, whereas depolymerization/deacetylation of tubulin inhibited IAV virion release ^59^. Furthermore, expression of HEV-ORF3 (HEV-1 Sar55 strain) in HeLa cells was discovered to induce tubulin acetylation ^19^. Therefore, we postulated that the ORF3-F10S mutation might affect the microtubule stability, and therefore examined the level of acetylated tubulin by Western blot. Like the ORF3 from HEV-1 Sar55 strain, the expression of HEV-3 ORF3 also enhanced tubulin acetylation (Figure 7B). However, the expression of F10S mutant resulted in dramatically reduced level of acetylated tubulin (Figure 7B), indicative of an influence of the F10S mutation on microtubule stability. To verify the role of F10S mutation in microtubule stability, we next examined the subcellular localization of Venus-tagged ORF3 and ORF3-F10S mutants. Consistent with previous report ^19^, Venus-tagged wild-type ORF3 showed a filamentous pattern of distribution and colocalized with tubulin in HeLa cells (Figure 7C). In contrast, the expression of Venus-tagged ORF3-F10S abolished the long filamentous structures in the cells (Figure 7C). In HEV-RNA transfected cells, enhanced tubulin acetylation was observed in S10-3 cells producing wild type HEV, compared with the MOCK transfected cells (1.75 VS 1) (Figure 7D). Although S10-3 cells producing the ORF3-F10S mutant virus did not show significant reduction in tubulin acetylation compared to the mock-treated cells (0.94 VS 1), the acetylated tubulin level detected in the mutant group was much lower than the cells transfected with wild-type HEV-RNA (0.94 VS 1.75, Figure 7D). These data demonstrate that the F10S mutation weakened the ability of the HEV-ORF3 protein to stabilize host microtubules during virus infection, and also suggest that prion-like aggregation of the ORF3 protein regulated the virion release process through modulating the microtubule stability.

### 3.7 Impaired prion aggregation of HEV-ORF3 causes reduced HEV pathogenesis *in vivo*

Given that the Mongolian gerbil can be infected by different HEV genotypes including HEV-3 KernowC1-p6 ^60–62^ we were able to use this animal model to explore the *in vivo* influence of HEV-ORF3 prion-like aggregation during HEV infection. Each gerbil was inoculated with wild type HEV or ORF3-F10S mutant at the dose of 1×10^9^ HEV genome copies. Based on the viremia level, a significant difference between the two groups of animal was observed at 1 week post infection. Gerbils infected by the wild type HEV produced much higher levels of viral RNA than gerbils infected by the ORF3-F10S mutant at 7 dpi (Figure 8A). The viremia disappeared in both groups after 14 dpi, which was consistent with previous observations that peak viremia in HEV-infected gerbils mainly lasts from 1 week to 2 weeks after virus inoculation. As HEV particles are largely shed through feces, HEV-RNA levels in fecal samples of the infected animals were examined as well. The results showed that animals infected by wild type HEV significantly shed more virus than those infected by the F10S mutant at 7 and 14 dpi, and that the fecal shedding of virus died down to similar levels in the two groups after 3 weeks of infection. Consistent with lower viremia levels and less fecal virus shedding, evaluation of anti-HEV-ORF2 antibodies in serum demonstrated a delayed serum conversion in gerbils infected by the ORF3-F10S mutant (Figure 8C). Compared with the serum conversion in all gerbils infected with wild type HEV at 21 dpi, only one gerbil was positive at 28 dpi and two out of five were positive at 35 and 42 dpi in the F10S group.

**Figure 8.**
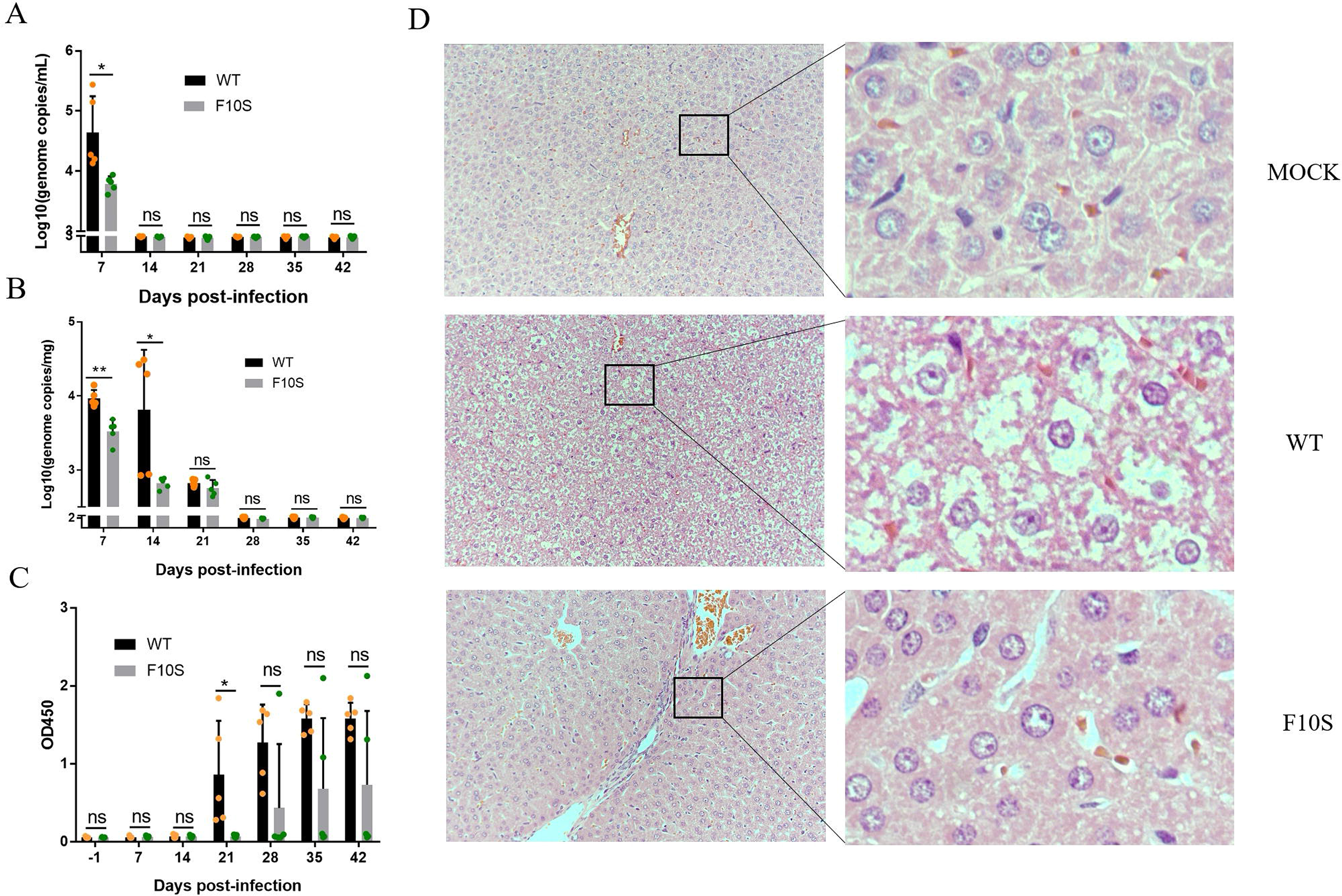
Mutation of HEV-ORF3 cPrD causes reduction of viral pathogenesis *in vivo*. **A.** Quantification of viral RNA in the sera of HEV infected Mongolian gerbils. The cDNA clone of HEV-3 KernowC1-p6 was used as reference for the calculation of HEV-RNA copies. **B**. Quantification of viral RNA in the fecal samples of HEV infected Mongolian gerbils. **C.** Examination of anti-HEV IgG levels in the sera of HEV infected Mongolian gerbils by ELISA. Recombinant ORF2-p239 of HEV-3 KernowC1-p6 was used as the coating antigen. **D.** Representative hematoxylin and eosin staining images of livers section of Mongolian gerbils infected by the indicated HEV at 14dpi.

As liver is the main target organ for HEV, livers were taken from infected gerbils at 2 weeks post infection and subjected to histological examination by hematoxylin and eosin (H&E) staining. Severe hydropic change of hepatocytes and congestion in liver section, the typical histological lesions in HEV infected gerbils ^61^, were observed in the wild type HEV infected gerbils (Figure 8D). In comparison, the pathological changes were much milder in the F10S group (Figure 8D). These data are consistent with the reduced viremia, less viral shedding and delayed serum conversion in the mutant virus-infected animals. Taken together, the *in vivo* animal study revealed that mutation of the ORF3 PrD to weaken its prion-like aggregation tendency also reduced the pathogenesis of the virus.

## 4. Discussion

Prions, originally defined as a portmanteau of pro (teinaceous) and in (fectious particle) ^63^, are self-propagating, transmissible protein particles. Prion-forming proteins can fold into multiple conformations are were initially discovered to be associated with neurodegenerative diseases in mammals, such as Kuru and Creutzfeldt–Jakob disease in human, scrapie in sheep and goat, and bovine spongiform encephalopathy in cattle ^64^. However, the discovery of prions in the microorganism *Saccharomyces cerevisiae* (yeast) and other microorganisms expanded the scope of the concept, and it is now accepted that prions are neither exclusive to mammals nor necessarily associated with disease. So far, prion-forming proteins have not only been identified in various cellular organisms including animal, plant ^38^, fungi ^65–68^ and bacteria ^39,69,70^, but also been discovered in a DNA virus ^35^. The prion-forming proteins so far described are structurally and functionally diverse, and the self-propagating properties of these proteins can serve as epigenetic information carriers that confer new traits to their hosts ^37,64^.

Although the first virus-encoded prion protein was only identified in a recent report ^35^, thousands of prion domains have been predicted by bioinformatic methods to be harbored in bacteriophages and eukaryotic viruses ^71,72^. In this study, we present our data that demonstrates that the HEV-ORF3 protein can form SDS-resistant high-molecular-weight aggregates in mammalian cells, and it is able to functionally replace the PrD of Sup35, a classical yeast prion, to display a [*PSI*^+^]-like phenotype in yeast. These features support that HEV-ORF3 is a virus-encoded prion protein and as such, is the first RNA virus encoded prion to be experimentally defined.

Similar to the prion-forming baculovirus LEF10 protein, the HEV-ORF3 protein has no Q/N-rich region which is commonly associated with amyloid-forming prions. Q/N-rich regions have been used as the principle selection criteria for the prediction of potential prion candidates by algorithms such as PLAAC ^56^. However, the first discovered and well-characterized prion protein, human PrP, also does not contain typical Q/N-rich regions. Other yeast prion-forming proteins linked to the [*ISP*^+^], [*MOD*^+^], [*GAR*^+^], [*ESI*^+^] and [*BIG*^+^] phenotypes are also atypical in this context ^73^. These atypical prions are poorly conserved in their linear amino acid sequence, but commonly contain intrinsically disordered regions (IDRs) which serve as PrDs to endow the proteins with prion-like behavior. The PrDs are often structurally independent and separable from the other regions in prion proteins. Here, our data demonstrate that the non-Q/N-rich ORF-3 protein and its cPrD are able to replace the typical Q/N-rich PrD of Sup35 to nucleate the conversion of Sup35 from a soluble monomer into aggregates and to confer it prion behavior thus supporting their role in prion conformational switching.

Microbial prions have been reported to play important regulatory roles in a wide variety of cellular processes and to confer various evolutionary advantages by forming self-propagation aggregates. In most cases, conversion of a protein into its inheritable prion form is associated with amyloid formation and believed to lead to a concomitant loss of protein function. In this study, we found that reduction of the prion-forming propensity of HEV-ORF3 disturbed its formation of filament-like structure, thereby weakening its ability to stabilize and harness host microtubules for virion release during virus replication. In line with its reduced ability to release progeny virus in cell culture, HEV-3 KernowC1-p6 bearing the F10S mutation demonstrated lower viremia, delayed serum conversion, and reduced viral pathogenesis during the infection of Mongolian Gerbils. These observations suggests that forming prion-like aggregates is important for HEV-ORF3 to serve its normal function.

Functional prion-like proteins have been described in multicellular organisms and unicellular microbes. For example, self-sustaining prion-like state of neuronal cytoplasmic polyadenylation element binding protein (CPEB) can be regulated by physiological signals and exploited for long-lasting memory ^74^ Microbial amyloids have been discovered that play important roles in surface-tension modulation, biofilm stabilization and adhesion ^75–77^. Our data identify HEV-ORF3 as a novel functional prion-like protein which assembles into filamentous structures for virus release, supporting the hypothesis that the beneficial forms of prions for the host harboring the gene could be functionally diverse and widespread in nature ^78^.

It has been reported that the HEV-ORF3 protein associates with the host cytoskeleton fraction and deletion of the N-terminal hydrophobic domain of ORF3 abolishes such association ^21^. However, co-IP data have also demonstrated that ORF3 does not directly interact with the microtubule components tubulin and microtubule-associated proteins (MAPs) ^19^. Visualization of microtubule filaments in IFA shows that ORF3 filaments are much less abundant than the microtubule filaments formed in the same cells. These observations imply that ORF3 filaments and microtubule filaments are related but not completely overlapped with each other and appear to be different. On the other hand, it has been reported that N-terminal PrD domain is required but not sufficient for microtubule association ^19^. Therefore, it appears that PrD mediated self-assembly of ORF3 is only one of the prerequisites for the formation of microtubule-associated filament structure. Further investigations are needed to elucidate the relationship between microtubules, the self-assembly characteristic and filament-like structure formed by ORF3, and the mechanism by which prion-like aggregation facilitates virion release.

In conclusion, we have discovered the first RNA virus-encoded prion protein, HEV-ORF3 and furthermore have demonstrated that the prion-like aggregation of the HEV-ORF3 protein is required to enhance microtubule stability in infected cells and then promote HEV virion release, which contributes to HEV pathogenesis in animal infection.

## Acknowledgement

This work was supported by a grant from the National Natural Science Foundation of China awarded to Y.N. (Grant No. 31672534), and a grant from Northwest A&F University awarded to H.C. (Grant No. Z10202190601).

## Author contributions

Y.N. and H.C. conceived the project, designed the research and supervised the experiments. Y.W., C.W., X.Z. and Q.Z. performed the mammalian cells and animal related experiments. H.T. and N.S. performed the yeast experiments. Y. H. provided technical guidance and support on the experiments on Mongolian gerbils. Y.Y. made the antibodies and contributed to the immuno-detection of HEV proteins. Q.D. contributed to the ORF3 mutants design and acquisition. M.T provided the yeast prion identification system. Y.N., H.C., Y.W., H. T. and M.T. analyzed the data and wrote the manuscript.

## Reference

1. Smith, D.B., Simmonds, P., Members Of The International Committee On The Taxonomy Of Viruses Study, G., Jameel, S., Emerson, S.U., Harrison, T.J., Meng, X.J., Okamoto, H., Van der Poel, W.H.M., and Purdy, M.A. (2014). Consensus proposals for classification of the family Hepeviridae. J Gen Virol 95, 2223–2232. 10.1099/vir.0.068429-0.

2. Nagashima, S., Takahashi, M., Kobayashi, T., Nishizawa, T., Nishiyama, T., Primadharsini, P.P., and Okamoto, H. (2017). Characterization of the Quasi-Enveloped Hepatitis E Virus Particles Released by the Cellular Exosomal Pathway. J Virol 91. 10.1128/JVI.00822-17.

3. Nan, Y., Wu, C., Zhao, Q., and Zhou, E.M. (2017). Zoonotic Hepatitis E Virus: An Ignored Risk for Public Health. Frontiers in microbiology 8, 2396. 10.3389/fmicb.2017.02396.

4. Jameel, S. (1999). Molecular biology and pathogenesis of hepatitis E virus. Expert Rev Mol Med 1999, 1–16. 10.1017/S1462399499001271.

5. Pillot, J., Turkoglu, S., Dubreuil, P., Cosson, A., Lemaigre, G., Meng, J., and Lazizi, Y. (1995). Cross-reactive immunity against different strains of the hepatitis E virus transferable by simian and human sera. C R Acad Sci III 318, 1059–1064.

6. Meng, X.J. (2013). Zoonotic and foodborne transmission of hepatitis E virus. Semin Liver Dis 33, 41–49. 10.1055/s-0033-1338113.

7. Dalton, H.R., Kamar, N., van Eijk, J.J., McLean, B.N., Cintas, P., Bendall, R.P., and Jacobs, B.C. (2016). Hepatitis E virus and neurological injury. Nature reviews. Neurology 12, 77–85. 10.1038/nrneurol.2015.234.

8. Geng, Y., Zhao, C., Huang, W., Harrison, T.J., Zhang, H., Geng, K., and Wang, Y. (2016). Detection and assessment of infectivity of hepatitis E virus in urine. J Hepatol 64, 37–43. 10.1016/j.jhep.2015.08.034.

9. Sadik, S., van Rijckevorsel, G.G., van Rooijen, M.S., Sonder, G.J., and Bruisten, S.M. (2016). Seroprevalence of hepatitis E virus differs in Dutch and first generation migrant populations in Amsterdam, the Netherlands: a cross-sectional study. BMC Infect Dis 16, 659. 10.1186/s12879-016-2007-z.

10. Feng, Z. (2016). Causation by HEV of extrahepatic manifestations remains unproven. Liver Int 36, 477–479. 10.1111/liv.13085.

11. Ahmad, I., Holla, R.P., and Jameel, S. (2011). Molecular virology of hepatitis E virus. Virus Res 161, 47–58. 10.1016/j.virusres.2011.02.011.

12. Tam, A.W., Smith, M.M., Guerra, M.E., Huang, C.C., Bradley, D.W., Fry, K.E., and Reyes, G.R. (1991). Hepatitis E virus (HEV): molecular cloning and sequencing of the full-length viral genome. Virology 185, 120–131. 10.1016/0042-6822(91)90760-9.

13. Tsarev, S.A., Emerson, S.U., Reyes, G.R., Tsareva, T.S., Legters, L.J., Malik, I.A., Iqbal, M., and Purcell, R.H. (1992). Characterization of a prototype strain of hepatitis E virus. Proc Natl Acad Sci U S A 89, 559–563. 10.1073/pnas.89.2.559.

14. Nair, V.P., Anang, S., Subramani, C., Madhvi, A., Bakshi, K., Srivastava, A., Shalimar, Nayak, B., Ranjith Kumar, C.T., and Surjit, M. (2016). Endoplasmic Reticulum Stress Induced Synthesis of a Novel Viral Factor Mediates Efficient Replication of Genotype-1 Hepatitis E Virus. PLoS Pathog 12, e1005521. 10.1371/journal.ppat.1005521.

15. Koonin, E.V., Gorbalenya, A.E., Purdy, M.A., Rozanov, M.N., Reyes, G.R., and Bradley, D.W. (1992). Computer-assisted assignment of functional domains in the nonstructural polyprotein of hepatitis E virus: delineation of an additional group of positive-strand RNA plant and animal viruses. Proc Natl Acad Sci U S A 89, 8259–8263. 10.1073/pnas.89.17.8259.

16. Graff, J., Torian, U., Nguyen, H., and Emerson, S.U. (2006). A bicistronic subgenomic mRNA encodes both the ORF2 and ORF3 proteins of hepatitis E virus. J Virol 80, 5919–5926. 10.1128/JVI.00046-06.

17. Mori, Y., and Matsuura, Y. (2011). Structure of hepatitis E viral particle. Virus Res 161, 59–64. 10.1016/j.virusres.2011.03.015.

18. Huang, Y.W., Opriessnig, T., Halbur, P.G., and Meng, X.J. (2007). Initiation at the third in-frame AUG codon of open reading frame 3 of the hepatitis E virus is essential for viral infectivity in vivo. J Virol 81, 3018–3026. 10.1128/JVI.02259-06.

19. Kannan, H., Fan, S., Patel, D., Bossis, I., and Zhang, Y.J. (2009). The hepatitis E virus open reading frame 3 product interacts with microtubules and interferes with their dynamics. J Virol 83, 6375–6382. 10.1128/JVI.02571-08.

20. Holla, R.P., Ahmad, I., Ahmad, Z., and Jameel, S. (2013). Molecular Virology of Hepatitis E Virus. Seminars in Liver Disease 33, 3–14. DOI 10.1055/s-0033-1338110.

21. Zafrullah, M., Ozdener, M.H., Panda, S.K., and Jameel, S. (1997). The ORF3 protein of hepatitis E virus is a phosphoprotein that associates with the cytoskeleton. J Virol 71, 9045–9053. 10.1128/JVI.71.12.9045-9053.1997.

22. Emerson, S.U., Nguyen, H., Torian, U., and Purcell, R.H. (2006). ORF3 protein of hepatitis E virus is not required for replication, virion assembly, or infection of hepatoma cells in vitro. J Virol 80, 10457–10464. 10.1128/JVI.00892-06.

23. Graff, J., Nguyen, H., Yu, C., Elkins, W.R., St Claire, M., Purcell, R.H., and Emerson, S.U. (2005). The open reading frame 3 gene of hepatitis E virus contains a cis-reactive element and encodes a protein required for infection of macaques. J Virol 79, 6680–6689. 10.1128/JVI.79.11.6680-6689.2005.

24. Yamada, K., Takahashi, M., Hoshino, Y., Takahashi, H., Ichiyama, K., Nagashima, S., Tanaka, T., and Okamoto, H. (2009). ORF3 protein of hepatitis E virus is essential for virion release from infected cells. J Gen Virol 90, 1880–1891. 10.1099/vir.0.010561-0.

25. Nagashima, S., Takahashi, M., Jirintai, Tanaka, T., Yamada, K., Nishizawa, T., and Okamoto, H. (2010). A PSAP motif in the ORF3 protein of hepatitis E virus is necessary for virion release from infected cells. Journal of General Virology 92, 269–278. 10.1099/vir.0.025791-0.

26. Nagashima, S., Takahashi, M., Jirintai, S., Tanaka, T., Nishizawa, T., Yasuda, J., and Okamoto, H. (2011). Tumour susceptibility gene 101 and the vacuolar protein sorting pathway are required for the release of hepatitis E virions. J Gen Virol 92, 2838–2848. 10.1099/vir.0.035378-0.

27. Ding, Q., Heller, B., Capuccino, J.M., Song, B., Nimgaonkar, I., Hrebikova, G., Contreras, J.E., and Ploss, A. (2017). Hepatitis E virus ORF3 is a functional ion channel required for release of infectious particles. Proc Natl Acad Sci U S A 114, 1147–1152. 10.1073/pnas.1614955114.

28. Nagashima, S., Takahashi, M., Jirintai, S., Tanggis, Kobayashi, T., Nishizawa, T., and Okamoto, H. (2014). The membrane on the surface of hepatitis E virus particles is derived from the intracellular membrane and contains trans-Golgi network protein 2. Arch Virol 159, 979–991. 10.1007/s00705-013-1912-3.

29. Hurley, J.H. (2010). The ESCRT complexes. Critical reviews in biochemistry and molecular biology 45, 463–487. 10.3109/10409238.2010.502516.

30. Feng, Z., Hirai-Yuki, A., McKnight, K.L., and Lemon, S.M. (2014). Naked Viruses That Aren't Always Naked: Quasi-Enveloped Agents of Acute Hepatitis. Annual review of virology 1, 539–560. 10.1146/annurev-virology-031413-085359.

31. Yin, X., Ambardekar, C., Lu, Y., and Feng, Z. (2016). Distinct Entry Mechanisms for Nonenveloped and Quasi-Enveloped Hepatitis E Viruses. J Virol 90, 4232–4242. 10.1128/JVI.02804-15.

32. Nan, Y., Ma, Z., Wang, R., Yu, Y., Kannan, H., Fredericksen, B., and Zhang, Y.J. (2014). Enhancement of interferon induction by ORF3 product of hepatitis E virus. J Virol 88, 8696–8705. 10.1128/JVI.01228-14.

33. Nan, Y., and Zhang, Y.J. (2016). Molecular Biology and Infection of Hepatitis E Virus. Frontiers in microbiology 7, 1419. 10.3389/fmicb.2016.01419.

34. Lhomme, S., Migueres, M., Abravanel, F., Marion, O., Kamar, N., and Izopet, J. (2020). Hepatitis E Virus: How It Escapes Host Innate Immunity. Vaccines 8. 10.3390/vaccines8030422.

35. Nan, H., Chen, H., Tuite, M.F., and Xu, X. (2019). A viral expression factor behaves as a prion. Nature communications 10, 359. 10.1038/s41467-018-08180-z.

36. Saupe, S.J. (2020). Amyloid Signaling in Filamentous Fungi and Bacteria. Annual review of microbiology 74, 673–691. 10.1146/annurev-micro-011320-013555.

37. Levkovich, S.A., Rencus-Lazar, S., Gazit, E., and Laor Bar-Yosef, D. (2021). Microbial Prions: Dawn of a New Era. Trends in biochemical sciences 46, 391–405. 10.1016/j.tibs.2020.12.006.

38. Chakrabortee, S., Kayatekin, C., Newby, G.A., Mendillo, M.L., Lancaster, A., and Lindquist, S. (2016). Luminidependens (LD) is an Arabidopsis protein with prion behavior. Proc Natl Acad Sci U S A 113, 6065–6070. 10.1073/pnas.1604478113.

39. Yuan, A.H., and Hochschild, A. (2017). A bacterial global regulator forms a prion. Science 355, 198–201. 10.1126/science.aai7776.

40. Emerson, S.U., Nguyen, H., Graff, J., Stephany, D.A., Brockington, A., and Purcell, R.H. (2004). In vitro replication of hepatitis E virus (HEV) genomes and of an HEV replicon expressing green fluorescent protein. J Virol 78, 4838–4846. 10.1128/jvi.78.9.4838-4846.2004.

41. Li, L., Xue, B., Sun, W., Gu, G., Hou, G., Zhang, L., Wu, C., Zhao, Q., Zhang, Y., Zhang, G., et al. (2018). Recombinant MYH9 protein C-terminal domain blocks porcine reproductive and respiratory syndrome virus internalization by direct interaction with viral glycoprotein 5. Antiviral Res 156, 10–20. 10.1016/j.antiviral.2018.06.001.

42. Gao, J., Xiao, S., Xiao, Y., Wang, X., Zhang, C., Zhao, Q., Nan, Y., Huang, B., Liu, H., and Liu, N.J.S.R. (2016). MYH9 is an Essential Factor for Porcine Reproductive and Respiratory Syndrome Virus Infection. 6, 25120.

43. Ju, X., Xiang, G., Gong, M., Yang, R., Qin, J., Li, Y., Nan, Y., Yang, Y., Zhang, Q.C., and Ding, Q. (2020). Identification of functional cis-acting RNA elements in the hepatitis E virus genome required for viral replication. PLoS Pathog 16, e1008488. 10.1371/journal.ppat.1008488.

44. Chen, Y., Liu, B., Sun, Y., Li, H., Du, T., Nan, Y., Hiscox, J.A., Zhou, E.M., and Zhao, Q. (2018). Characterization of Three Novel Linear Neutralizing B-Cell Epitopes in the Capsid Protein of Swine Hepatitis E Virus. J Virol 92. 10.1128/JVI.00251-18.

45. Yang, Y., Lin, S., Nan, Y., Ma, Z., Yang, L., and Zhang, Y. (2016). A Linear Surface Epitope in a Proline-Rich Region of ORF3 Product of Genotype 1 Hepatitis E Virus. Viruses 8. 10.3390/v8080227.

46. Gietz, R.D., Schiestl, R.H., Willems, A.R., and Woods, R.A. (1995). Studies on the transformation of intact yeast cells by the LiAc/SS-DNA/PEG procedure. Yeast 11, 355–360. 10.1002/yea.320110408.

47. Patel, D., Opriessnig, T., Stein, D.A., Halbur, P.G., Meng, X.J., Iversen, P.L., and Zhang, Y.J. (2008). Peptide-conjugated morpholino oligomers inhibit porcine reproductive and respiratory syndrome virus replication. Antiviral Res 77, 95–107. 10.1016/j.antiviral.2007.09.002.

48. Halfmann, R., and Lindquist, S. (2008). Screening for amyloid aggregation by Semi-Denaturing Detergent-Agarose Gel Electrophoresis. Journal of visualized experiments: JoVE. 10.3791/838.

49. Alberti, S., Halfmann, R., King, O., Kapila, A., and Lindquist, S. (2009). A systematic survey identifies prions and illuminates sequence features of prionogenic proteins. Cell 137, 146–158. 10.1016/j.cell.2009.02.044.

50. Chernoff, Y.O., Lindquist, S.L., Ono, B., Inge-Vechtomov, S.G., and Liebman, S.W. (1995). Role of the chaperone protein Hsp104 in propagation of the yeast prion-like factor [psi+]. Science 268, 880–884. 10.1126/science.7754373.

51. Jung, G., and Masison, D.C. (2001). Guanidine hydrochloride inhibits Hsp104 activity in vivo: a possible explanation for its effect in curing yeast prions. Current microbiology 43, 7–10. 10.1007/s002840010251.

52. Shorter, J., and Lindquist, S. (2004). Hsp104 catalyzes formation and elimination of self-replicating Sup35 prion conformers. Science 304, 1793–1797. 10.1126/science.1098007.

53. Nan, Y., Wu, C., Zhao, Q., Sun, Y., Zhang, Y.J., and Zhou, E.M. (2018). Vaccine Development against Zoonotic Hepatitis E Virus: Open Questions and Remaining Challenges. Frontiers in microbiology 9, 266. 10.3389/fmicb.2018.00266.

54. Conchillo-Sole, O., de Groot, N.S., Aviles, F.X., Vendrell, J., Daura, X., and Ventura, S. (2007). AGGRESCAN: a server for the prediction and evaluation of "hot spots" of aggregation in polypeptides. BMC bioinformatics 8, 65. 10.1186/1471-2105-8-65.

55. Garbuzynskiy, S.O., Lobanov, M.Y., and Galzitskaya, O.V. (2010). FoldAmyloid: a method of prediction of amyloidogenic regions from protein sequence. Bioinformatics 26, 326–332. 10.1093/bioinformatics/btp691.

56. Lancaster, A.K., Nutter-Upham, A., Lindquist, S., and King, O.D. (2014). PLAAC: a web and command-line application to identify proteins with prion-like amino acid composition. Bioinformatics 30, 2501–2502. 10.1093/bioinformatics/btu310.

57. Yin, X., Ying, D., Lhomme, S., Tang, Z., Walker, C.M., Xia, N., Zheng, Z., and Feng, Z. (2018). Origin, antigenicity, and function of a secreted form of ORF2 in hepatitis E virus infection. Proc Natl Acad Sci U S A 115, 4773–4778. 10.1073/pnas.1721345115.

58. Westermann, S., and Weber, K. (2003). Post-translational modifications regulate microtubule function. Nature reviews 4, 938–947. 10.1038/nrm1260.

59. Husain, M., and Harrod, K.S. (2011). Enhanced acetylation of alpha-tubulin in influenza A virus infected epithelial cells. FEBS Lett 585, 128–132. 10.1016/j.febslet.2010.11.023.

60. Li, W., Sun, Q., She, R., Wang, D., Duan, X., Yin, J., and Ding, Y. (2009). Experimental infection of Mongolian gerbils by a genotype 4 strain of swine hepatitis E virus. J Med Virol 81, 1591–1596. 10.1002/jmv.21573.

61. Xu, L.D., Zhang, F., Chen, C., Peng, L., Luo, W.T., Chen, R., Xu, P., and Huang, Y.W. (2022). Revisiting the Mongolian Gerbil Model for Hepatitis E Virus by Reverse Genetics. Microbiology spectrum 10, e0219321. 10.1128/spectrum.02193-21.

62. Zhang, W., Ami, Y., Suzaki, Y., Doan, Y.H., Muramatsu, M., and Li, T.C. (2022). Mongolia Gerbils Are Broadly Susceptible to Hepatitis E Virus. Viruses 14. 10.3390/v14061125.

63. Bolton, D.C., McKinley, M.P., and Prusiner, S.B. (1982). Identification of a protein that purifies with the scrapie prion. Science 218, 1309–1311. 10.1126/science.6815801.

64. Scheckel, C., and Aguzzi, A. (2018). Prions, prionoids and protein misfolding disorders. Nature reviews. Genetics 19, 405–418. 10.1038/s41576-018-0011-4.

65. Wickner, R.B. (1994). [URE3] as an altered URE2 protein: evidence for a prion analog in Saccharomyces cerevisiae. Science 264, 566–569. 10.1126/science.7909170.

66. Patel, B.K., Gavin-Smyth, J., and Liebman, S.W. (2009). The yeast global transcriptional co-repressor protein Cyc8 can propagate as a prion. Nat Cell Biol 11, 344–349. 10.1038/ncb1843.

67. Du, Z., Park, K.W., Yu, H., Fan, Q., and Li, L. (2008). Newly identified prion linked to the chromatin-remodeling factor Swi1 in Saccharomyces cerevisiae. Nature genetics 40, 460–465. 10.1038/ng.112.

68. Coustou, V., Deleu, C., Saupe, S., and Begueret, J. (1997). The protein product of the het-s heterokaryon incompatibility gene of the fungus Podospora anserina behaves as a prion analog. Proc Natl Acad Sci U S A 94, 9773–9778. 10.1073/pnas.94.18.9773.

69. Shahnawaz, M., Park, K.W., Mukherjee, A., Diaz-Espinoza, R., and Soto, C. (2017). Prion-like characteristics of the bacterial protein Microcin E492. Scientific reports 7, 45720. 10.1038/srep45720.

70. Fleming, E., Yuan, A.H., Heller, D.M., and Hochschild, A. (2019). A bacteria-based genetic assay detects prion formation. Proc Natl Acad Sci U S A 116, 4605–4610. 10.1073/pnas.1817711116.

71. Tetz, G., and Tetz, V. (2017). Prion-Like Domains in Phagobiota. Frontiers in microbiology 8, 2239. 10.3389/fmicb.2017.02239.

72. Tetz, G., and Tetz, V. (2018). Prion-like Domains in Eukaryotic Viruses. Scientific reports 8, 8931. 10.1038/s41598-018-27256-w.

73. Dennis, E.M., and Garcia, D.M. (2022). Biochemical Principles in Prion-Based Inheritance. Epigenomes 6. 10.3390/epigenomes6010004.

74. Majumdar, A., Cesario, W.C., White-Grindley, E., Jiang, H., Ren, F., Khan, M.R., Li, L., Choi, E.M., Kannan, K., Guo, F., et al. (2012). Critical role of amyloid-like oligomers of Drosophila Orb2 in the persistence of memory. Cell 148, 515–529. 10.1016/j.cell.2012.01.004.

75. Wosten, H.A., and de Vocht, M.L. (2000). Hydrophobins, the fungal coat unravelled. Biochimica et biophysica acta 1469, 79–86. 10.1016/s0304-4157(00)00002-2.

76. Chapman, M.R., Robinson, L.S., Pinkner, J.S., Roth, R., Heuser, J., Hammar, M., Normark, S., and Hultgren, S.J. (2002). Role of Escherichia coli curli operons in directing amyloid fiber formation. Science 295, 851–855. 10.1126/science.1067484.

77. Ramsook, C.B., Tan, C., Garcia, M.C., Fung, R., Soybelman, G., Henry, R., Litewka, A., O'Meally, S., Otoo, H.N., Khalaf, R.A., et al. (2010). Yeast cell adhesion molecules have functional amyloid-forming sequences. Eukaryot Cell 9, 393–404. 10.1128/EC.00068-09.

78. Si, K. (2015). Prions: what are they good for? Annual review of cell and developmental biology 31, 149–169. 10.1146/annurev-cellbio-100913-013409.

