## Supplementary Table 1 for "Prion-like characteristics of Hepatitis E virus ORF3 protein are associated with virus release and pathogenesis"

**Supplementary Table 1.** List of primers and corresponding sequence used

| Primers | Sequence (5'-3') | Description |
| --- | --- | --- |
| JVHEV-F | GGTGGTTTCTGGGGTGAC | Primer pairs for Taqman probe |
| JVHEV-R | AGGGGTTGGTTGGATGAA |  |
| JVHEVP | TGATTCTCAGCCCTTCGC | Taqman Probe |
| ORF1-F | AATCCAGACCACGAGCCG | qPCR for detection of HEV(+)/(-) RNA |
| ORF1-R | ATCGCACCAGGGTTAGCG |  |
| VN173-ORF3-F | CGGGATCCAtgggttcgcgaccatgcgcc |  |
| VN173-ORF3-R | GCAAGCTTACTGCCTCCACCGCCACTGCCTCCACCGCCgcggcgcggccccagctgtg |  |
| VC155-ORF3-F | CGGGATCCAtgggttcgcgaccatgcgccc |  |
| VC155-ORF3-R | AAGCTTACTGCCTCCACCGCCACTGCCTCCACCGCCgcggcgcggccccagct |  |
| PCAGEN-GT-3-ORF3-F | CGgaattcatgggatcaccatgtgccctagg |  |
| PCAGEN-GT-3-ORF3-R | GCctcgagtcaacggcgcagccccagctgg |  |
| PCAGEN-GT-1-ORF3-F | GCgaattcatgggttcgcgaccatg |  |
| PCAGEN-GT-1-ORF3-R | GCctcgaggcggcgcggccccagctgtggta |  |
| PCAGEN-GT-2-ORF3-F | GGgaattcAtgggttcgccaccatgcg |  |
| PCAGEN-GT-2-ORF3-R | GGctcgagtcaCAGATCCTCTTCAGAGATGAG |  |
| PCAGEN-GT-4-ORF3-F | GGgaattcatggagatgccaccatgcgc |  |
| PCAGEN-GT-4-ORF3-R | GGctcgagacggcgaagccccagctggggcag |  |
| VC155-ORF3-N25-F | CGGGATCCAtgggttcgcgaccatgcgcc |  |
| VC155-ORF3-N25-R | GCAAGCTTACTGCCTCCACCGCCACTGCCTCCACCGCCgtggcgcgggcagcataggcag |  |
| VC155-ORF3-C90-F | CGGGATCCAtgcgcccggtcagccgtctg |  |
| VC155-ORF3-C90-R | GCAAGCTTACTGCCTCCACCGCCACTGCCTCCACCGCCgcggcgcggccccagctgtgg |  |
| VN173-ORF3-N25-F | CGGGATCCAtgggttcgcgaccatgcgcc |  |
| VN173-ORF3-N25-R | AAGCTTACTGCCTCCACCGCCACTGCCTCCACCGCCgtggcgcgggcagcataggcag |  |
| VN173-ORF3-C90-F | CGGGATCCAtgcgcccggtcagccgtctg |  |
| VN173-ORF3-C90-R | ctaccacagctggggccgcgccgcGGCGGTGGAGGCAGTGGCGGTGGAGGCAGTAAGCTT |  |
| VN173-GT-1-ORF3-N25-F | GCgaattcatgggttcgcgaccatg |  |
| VN173-GT-1-ORF3-N25-R | CAAGCTTACTGCCTCCACCGCCACTGCCTCCACCGCCccggtggcgcgggcagcat |  |
| VN173-GT-2-ORF3-N25-F | CGGATCCAtgggttcgccacca |  |
| VN173-GT-2-ORF3-N25-R | CAAGCTTACTGCCTCCACCGCCACTGCCTCCACCGCCgtggcgcgggcaacatagg |  |
| VN173-GT-4-ORF3-N25-F | CGGATCCatggagatgccaccat |  |
| VN173-GT-4-ORF3-N25-R | CAAGCTTACTGCCTCCACCGCCACTGCCTCCACCGCCgtggcgcgggcagcata |  |
| PCAGEN-ORF3-M1-F | GGgaattcatggccgccgcctgtgccctaggg |  |
| PCAGEN-ORF3-M1-R | GGctcgagtcaacggcgcagccccagctgggg |  |
| PCAGEN-ORF3-M2-F | GGgaattcatgggatcaccagccgccgccggg |  |
| PCAGEN-ORF3-M2-R | GGctcgagtcaacggcgcagccccagctgggg |  |
| PCAGEN-ORF3-M3-F | GGgaattcatgggatcaccatgtgccctagccgccgcctgct |  |
| PCAGEN-ORF3-M3-R | GGctcgagtcaacggcgcagccccagctgggg |  |
| PCAGEN-ORF3-M4-F | GGgaattcatgggatcaccatgtgccctagggttgttcgccgccgcctcttcgt |  |
| PCAGEN-ORF3-M4-R | GGctcgagtcaacggcgcagccccagctgggg |  |
| PCAGEN-ORF3-M5-F | GGgaattcatgggatcaccatgtgccctagggttgttctgctgctgtgccgccgccttctgc |  |
| PCAGEN-ORF3-M5-R | GGctcgagtcaacggcgcagccccagctgggg |  |
| PCAGEN-ORF3-M6-F | GGgaattcatgggatcaccatgtgccctagggttgttctgctgctgttcttcgtgtgccgccgcctgctgcccg |  |
| PCAGEN-ORF3-M6-R | GGctcgagtcaacggcgcagccccagctgggg |  |
| PCAGEN-ORF3-M7-F | GGgaattcatgggatcaccatgtgccctagggttgttctgctgctgttcttcgtgtttctgcctagccgccgcccgcca |  |
| PCAGEN-ORF3-M7-R | GGctcgagtcaacggcgcagccccagctgggg |  |
| PCAGEN-ORF3-M8-F | GGgaattcatgggatcaccatgtgccctagggttgttctgctgctgttcttcgtgtttctgcctatgctgcccggccgccgccccgg |  |
| PCAGEN-ORF3-M8-R | GGctcgagtcaacggcgcagccccagctgggg |  |
| PCAGEN-ORF3-F8A-F | gGGaattcatgggatcaccatgtgccctagccttgttctgctgc |  |
| PCAGEN-ORF3-F8A-R | GGctcgagtcaacggcgcagccccagctgggg |  |
| PCAGEN-ORF3-F9A-F | gGGaattcatgggatcaccatgtgccctaggggccttctgctgc |  |
| PCAGEN-ORF3-F9A-R | GGctcgagtcaacggcgcagccccagctgggg |  |
| PCAGEN-ORF3-F10A-F | gGGaattcatgggatcaccatgtgccctagggttggcctgctgctgttcttcg |  |
| PCAGEN-ORF3-F10A-R | GGctcgagtcaacggcgcagccccagctgggg |  |
| PCAGEN-ORF3-F10C-F | gGGaattcatgggatcaccatgtgccctagggttgtgttgctgctgttc |  |
| PCAGEN-ORF3-F10C-R | GGctcgagtcaacggcgcagccccagctgggg |  |
| PCAGEN-ORF3-F10S-F | gGGaattcatgggatcaccatgtgccctagggttgtcttgctgctg |  |
| PCAGEN-ORF3-F10S-R | GGctcgagtcaacggcgcagccccagctgggg |  |
| PCAGEN-ORF3-F10Y-F | gGGaattcatgggatcaccatgtgccctagggttgtattgctgctgttc |  |
| PCAGEN-ORF3-F10Y-R | GGctcgagtcaacggcgcagccccagctgggg |  |
| KernowC1-p6-ORF3-F1 | gtgccctagggttgtcttgctgctgttc |  |
| KernowC1-p6-ORF3-R1 | ACagcccctgtacctgatgttgattcacgtg |  |
| KernowC1-p6-ORF3-F2 | CTcttaagggtttctggaagaagcat |  |
| KernowC1-p6-ORF3-R2 | gtgccctagggttgtcttgctgctgttc |  |
| KernowC1-p6-F | ataaccttattggcatgttgcagaccat |  |
| Sup35M-F | cactcgaccaaagctcccattgc |  |
| Sup35M-R | ttcttatcagattcggcagg |  |
| Sar55-F | tgctctagaggttcgcgaccatgcgcc |  |
| Sar55-R | ccggaattcgcggcgcggccccagctg |  |
| Sar551-25F | aactctagaggttcgcgaccatg |  |
| Sar551-25R | tctgaattcgcggtggcgcgggcagca |  |
| Sar5526-113F | tcttctagaccggtcagccgtctggcc |  |
| Sar5526-113R | catgaattcgcggcgcgg |  |
| MEX-F | tgctctagaggttcgccaccatgcgccct |  |
| MEX-R | ccggaattcgcgccgcagccccggctgtg |  |
| MEX1-25F | aactctagaggttcgccaccatg |  |
| MEX1-25R | tctgaattctcggtggcgcgggcaac |  |
| MEX26-113F | tcttctagaccggtcagccgtctgg |  |
| MEX26-113R | catgaattcgcgccgcagc |  |
| Ker-F | tgctctagaggatcaccatgtgccctagg |  |
| Ker-R | ccggaattcacggcgcagccccagctggg |  |
| Ker1-25 -F | tgctctagaggatcaccatgtgccctagg |  |
| Ker1-25 -R | ccggaattcccggtggcgcgggcagcatag |  |
| Ker26-113 -F | tgctctagaccggccagccgtctggc |  |
| Ker26-113 -R | ccggaattcacggcgcagccccagctggg |  |
| Ker1-60 -F | tgctctagaggatcaccatgtgccctagg |  |
| Ker1-60 -R | ccggaattcgaatataggggagggcgaag |  |
| Ker26-60 -F | tgctctagaccggccagccgtctggc |  |
| Ker26-60 -R | ccggaattcgaatataggggagggcgaag |  |
| Ker61-113 -F | tgctctagaatccaaccaaccccttcgcc |  |
| Ker61-113 -R | ccggaattcacggcgcagccccagctggg |  |
| SD-F | tgctctagagagatgccaccatgcgctct |  |
| SD-R | ccggaattcacggcgaagccccagctggg |  |
| SD1-25 -F | aactctagagagatgccaccatgc |  |
| SD1-25 -R | tctgaattcccggtggcgcgggcagcata |  |
| SD26-113 -F | tcttctagaccggtcagccgtctggc |  |
| SD26-113 -R | catgaattcacggcgaagccc |  |
| HSP104-Del-F | GGCAAAGGGGCGCAAACTTATGCAACCTGCCAGATTATTATATAAGGCgagcagattgtactgagagtgcacc |  |
| HSP104-Del-R | CAATTTCCATACTGTCCTCATTATCGTCATCACCTAACGTGTCAGCCttagttttgctggccgcatcttctc |  |
| Del-test-F | attgaaccctccatcgtggtag |  |
| hsp104-test-R | ggaacaagtgacaaaggaacga |  |
| URA3-test-R | aattgtacttggcggataatgc |  |
| Sar55-SacⅡ-F | tccccgcggatgggttcgcgaccatgc |  |
| Sar55-NheⅠ-R | ctagctagcgcggcgcggccccagctgtgg |  |
| Ker-SacⅡ-F | tccccgcggatgggatcaccatgtgcc |  |
| Ker-NheⅠ-R | ctagctagcacggcgcagccccagctg |  |
| KerM1-F | tgctctagagccgccgcctgtgccctagg |  |
| KerM2-F | tgctctagaggatcaccagccgccgccgg |  |
| KerM1- SacⅡ-F | tccccgcggatggccgccgcctgtgcc |  |
| KerM2- SacⅡ-F | tccccgcggatgggatcaccagccgcc |  |
