## Supplementary figures and images for "Prion-like characteristics of Hepatitis E virus ORF3 protein are associated with virus release and pathogenesis"

### Supplementary Figure 1

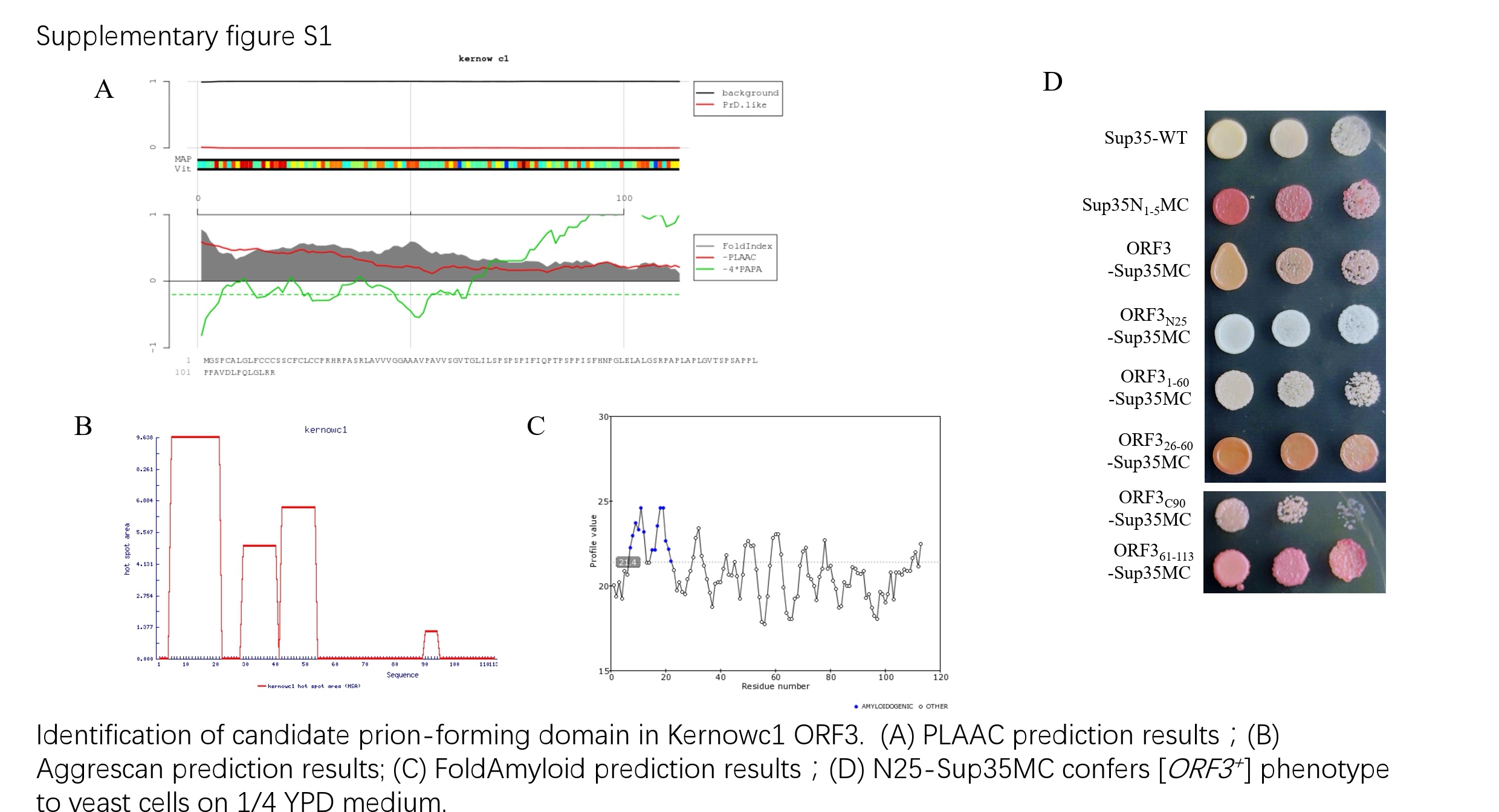

### Supplementary Figure 2

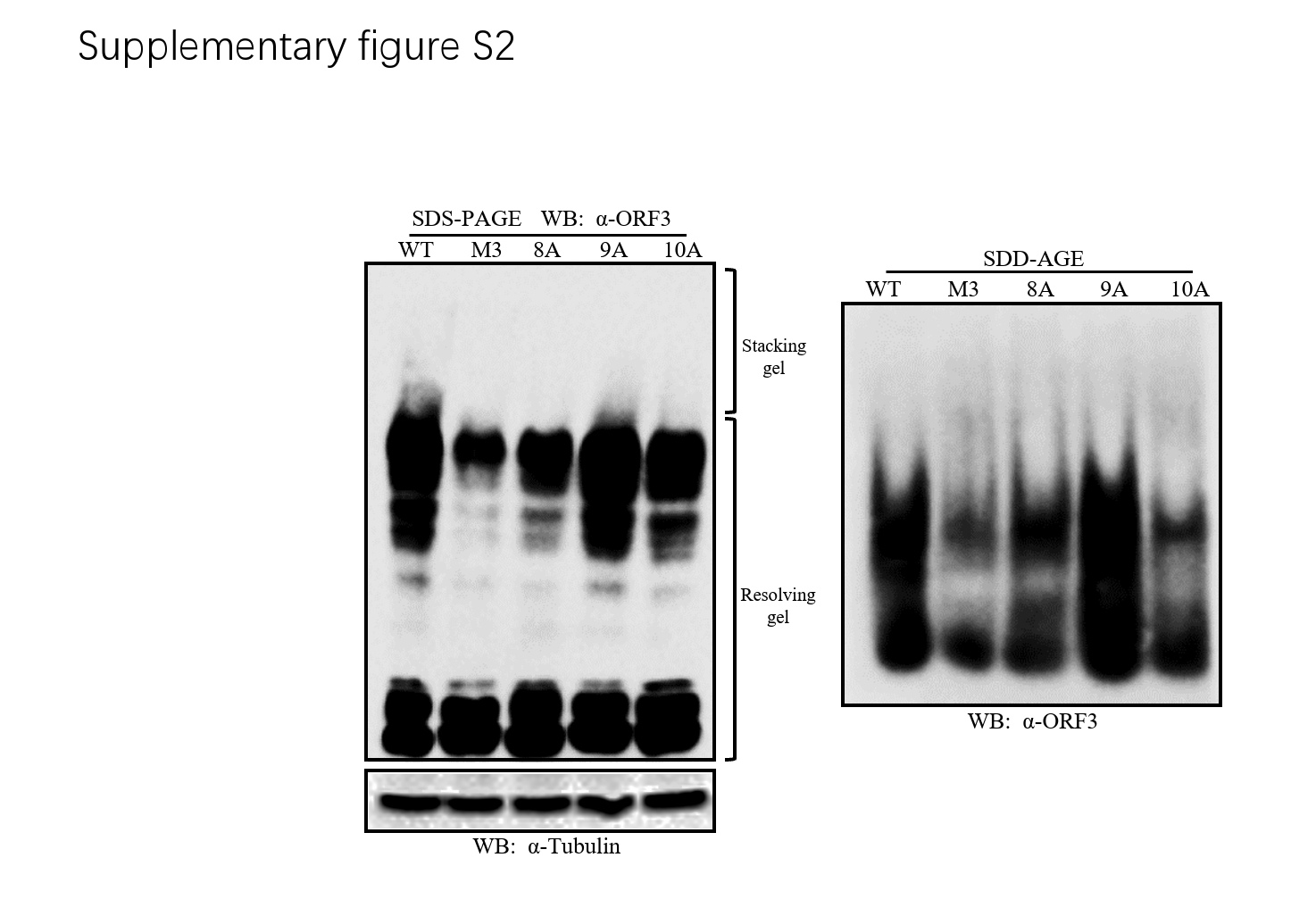
